# Regulation of Female Reproductive Aging by the *Spag17* Gene

**DOI:** 10.1101/2025.03.01.640648

**Authors:** Valerie Ericsson, Madisyn Elam, Paulene Sapao, Le My Tu Nguyen, Mary Ellen Gill, Saikeerthana Chodavarapu, Carlos Córdova-Fletes, Audrey Lafrenaye, Sonia Hassan, Mala Mahendroo, Li Chen, Mathieu Petitjean, Jerome F Strauss, John Varga, Francesca E Duncan, Maria E Teves

## Abstract

Reproductive aging in females is characterized by a decline in oocyte quantity and quality, as well as uterine and cervical dysfunction that contributes to infertility and pregnancy complications. To investigate mechanisms underlying reproductive aging, we explored the contribution of *Spag17*, a cilia-related gene associated with tissue homeostasis and fibrosis. *Spag17* was expressed throughout the female reproductive tract; however, its expression declined with age in ovarian tissue, while high expression levels were observed in the cervix of young females during cervical tissue remodeling in the pre- and post-parturition periods. Loss of *Spag17* in mice resulted in impaired fertility, obstructed labor, and maternal death. This phenotype was associated with accelerated ovarian aging, increased fibrosis, and cervical stiffness, further complicating parturition. At the molecular level, *Spag17* loss activated key aging-associated pathways, including proinflammatory, profibrotic, and senescence signaling, suggesting that SPAG17 may be a critical player in female reproductive aging.

**TEASER:** *Spag17* is a key modulator of female reproductive aging.

## INTRODUCTION

Aging is a major determinant of female reproductive well-being and one of the biggest challenges in reproductive medicine for which there are no clinically proven therapies. Advanced female age contributes to infertility, pregnancy complications, and congenital defects in the offspring [Duncan et al., 2018]. With the global rise in mean maternal age, there is a pressing need to prevent age-associated infertility and adverse pregnancy outcomes.

The organs of the female reproductive tract exhibit significant susceptibility to age-related pathologies [Balough et al., 2024]. In the uterus, the most notable signs of aging include endometrial atrophy, uterine fibrosis, decreased vascular compliance, and diminished uterine vascular adaptation to pregnancy [Wu et al., 2023]. Aging is also accompanied by significant changes in cell-type composition. For example, the uterus shows a reduction in fibroblasts and an increase in M1 and M2 macrophages; the cervix and vagina exhibit a decrease in M1 macrophages; the oviduct demonstrates an increase in NK, B, and dendritic cells; and the ovaries show a decrease in follicle-associated cells such as theca cells, granulosa cells, and luteal cells, along with an increase in fibroblasts and immune cells [Winkler et al., 2024].

Moreover, ovarian aging is characterized by the excessive accumulation of extracellular matrix, which replaces parenchymal tissue in the ovarian stroma [Briley et al., 2016; Dipali et al., 2023], accompanied by increased stiffness, fibrosis, and lower expression of hyaluronan [Amargant et al., 2020; Umehara et al., 2018]. Additionally, higher levels of inflammatory cytokines [Briley et al., 2016] and inflammatory gene expression [Zhang et al., 2020] are associated with an increased presence of multinucleated macrophage giant cells (MNGCs) [Briley et al., 2016; Foley et al., 2021] and other immune cell populations [Isola et al., 2024; Winkler et al., 2024]. At the transcriptional level, increased expression of proinflammatory, profibrotic, and senescence markers has been observed in aged mouse ovaries [Isola et al., 2024].

Although aging occurs at varying rates in different organ systems, the aging process can be attributed to cellular senescence. Senescent cells, as a consequence of arrest in the cell cycle, secrete inflammatory cytokines, interleukins, and growth factors that influence the behaviors of surrounding cells. This is described as the “senescence associated secretory phenotype” (SASP) [Almeida et al, 2017; Suryadevara et al., 2024]. Moreover, senescent fibroblasts release chemokines and proteases that target the extracellular matrix (ECM) and promote fibrosis, thereby affecting the tissue structure [Davalos et al., 2010].

Our recent investigation has uncovered a novel mechanistic pathway linking aging and fibrosis to the Sperm-associated antigen 17 (*Spag17*) gene [Sapao et al., 2023]. *Spag17* is a pleiotropic gene expressed in a wide variety of cell types. While SPAG17 protein is primarily recognized for its role in motile cilia/flagella [Teves et al., 2013], emerging evidence demonstrates its essential functions in other cell types and tissues [Teves et al., 2014 and 2015; Kazarian et al., 2018; Agudo-Rios et al., 2023]. Notably, recent studies have shown that the loss of SPAG17 promotes the transition of fibroblasts into myofibroblasts, leading to disrupted tissue remodeling and fibrosis in the skin and other organs [Sapao et al., 2018; Sapao et al., 2023]. Furthermore, SPAG17 deficiency in dermal fibroblasts has been associated with differential expression of genes involved in extracellular matrix organization and senescence-related mechanisms [Sapao et al., 2023]. These findings suggest that SPAG17 may act as a key regulator of cellular processes associated with tissue aging and fibrosis across organs.

Given this evidence, we explore the role of *Spag17* in reproductive function. Our studies revealed that SPAG17 deficiency adversely affects ovarian, and cervical functions, ultimately leading to reduced fertility and parturition defects. These reproductive phenotypes are likely driven by age-related tissue remodeling and fibrotic processes mediated by SPAG17 dysregulation. Collectively, these findings highlight SPAG17 as a critical regulator of aging and fibrosis in the female reproductive tract. This novel insight underscores the importance of SPAG17 in maintaining reproductive tissue homeostasis and provides a molecular target for future therapeutic strategies to improve age-related reproductive decline and fibrotic disorders.

## RESULTS

### *SPAG17* expression in the female reproductive tract

Understanding the role of SPAG17 in the female reproductive tract is crucial given its involvement in tissue remodeling, aging, and fibrosis. To explore its impact, we investigated *SPAG17* expression in both human and mouse reproductive tissues using a combination of publicly available genomic data, immunohistochemistry and β-Galactosidase staining.

Using the GTEx Portal, we identified *SPAG17* mRNA expression across the entire female reproductive tract. Notably, *SPAG17* expression was highest in the fallopian tube, cervix, and vagina, with comparatively lower levels detected in the uterus and ovary (Fig. 1A). Single-cell RNA sequencing (scRNAseq) studies provided further insight into the cellular distribution of *SPAG17*. In the mouse cervix, data from Guo and collaborators [Guo et al., 2022] revealed *Spag17* expression predominantly in fibroblasts, stromal cells, endothelial and smooth muscle cells, with lower expression in immune cells (Fig. 1B).

**Figure 1:**
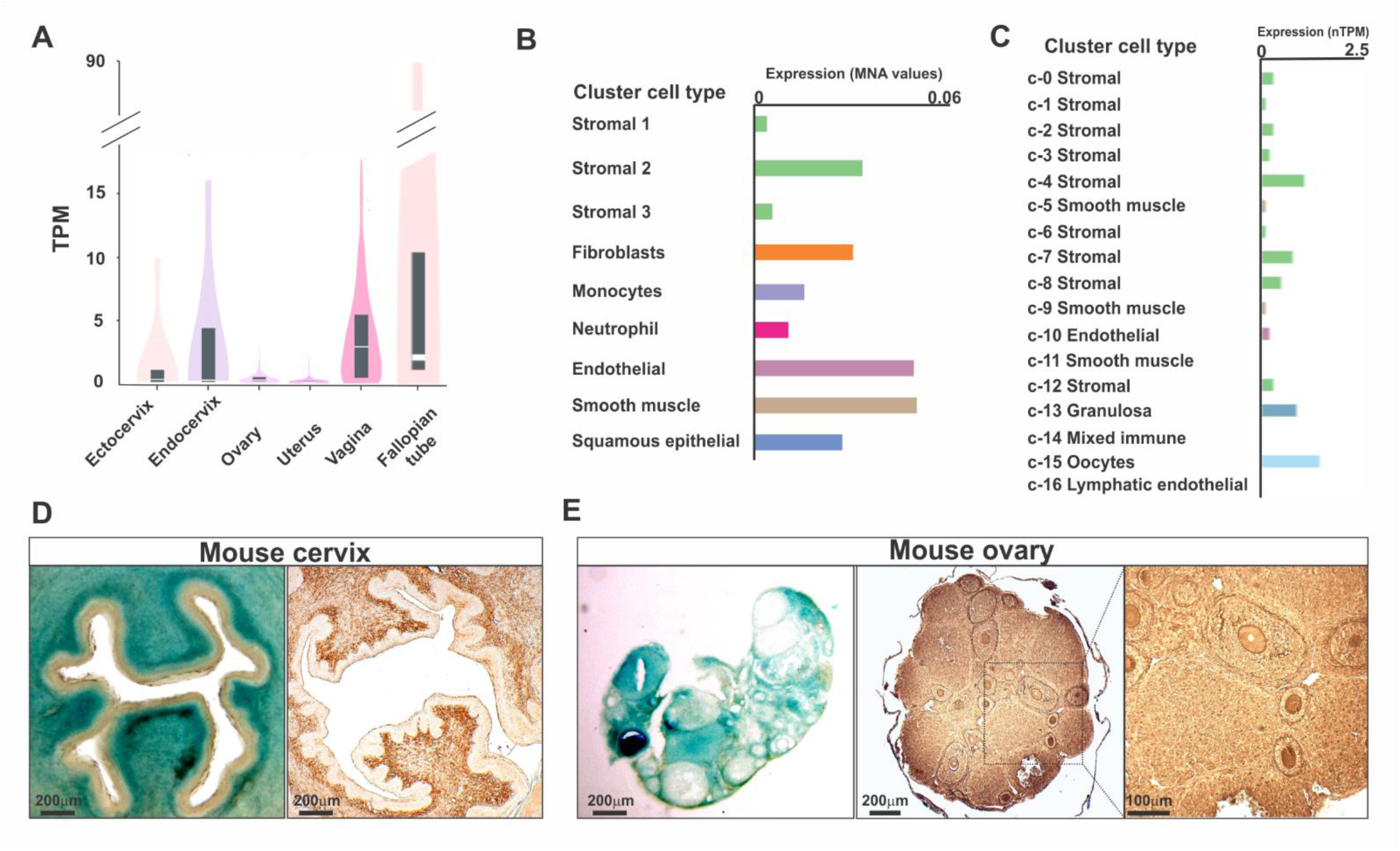
Spag17 expression in the female reproductive tract. (**A**) Half violin plot of expression patterns of *SPAG17* mRNA across different female reproductive tissues obtained from GTExPortal. The Y-axis indicates TPM (transcripts per million). Bar charts of *SPAG17* scRNAseq expression in various cell type clusters within the mouse cervix (**B**) and human ovary (**C**). Representative images of β-galactosidase staining (n=3) and immunohistochemistry (n=6) detection of SPAG17 in mouse cervix (**D**) and mouse ovary (**E**). Scale bars, 100 µm and 200 µm.

In ovarian tissue, data from the Human Protein Atlas (proteinatlas.org) revealed that *SPAG17* expression is highest in stromal cells, granulosa/theca cells, and oocytes, while smooth muscle and endothelial cells exhibited the lowest expression levels (Fig. 1C).

These findings underscore significant expression of *SPAG17* in key cell types of the cervix and ovary, suggesting a pivotal role in maintaining the structure and function of these tissues.

To validate this in silico analysis, we conducted further experimental studies to evaluate SPAG17 expression in the mouse cervix and ovary. β-Galactosidase staining was performed on *Spag17/Sox2-Cre* heterozygous mice (n=3), which possess an insertion tagging the *Spag17* gene with a *LacZ* reporter, to determine the prevalence and localization of *Spag17* expression in the cervix and ovaries. Intense blue staining was observed in the stroma of cervical tissue (Fig. 1D; Fig. S1A), consistent with results obtained by immunohistochemistry (n=6) using an anti-SPAG17 antibody (Fig. 1D; Fig. S2A).

In the ovaries, β-galactosidase staining was distributed among oocytes, granulosa cells, stromal cells, theca cells, and corpora lutea (n=3; Fig. 1E; Fig. S1B and C). Similar results were observed by immunohistochemistry in wild-type mouse ovaries (n=6; Fig. 1E), whereas no immunohistochemical staining was detected in the negative controls or in samples from *Spag17* knockout mice (Fig. S2B). Overall, the observed SPAG17 distribution in the mouse was consistent with the in silico genomic data.

### Loss of *SPAG17* results in reduced fertility and pregnancy complications

To investigate the role of SPAG17 in female fertility we used the *Spag17/Sox2-Cre* knockout mouse model [Kazarian et al., 2018]. Reproductive-age wild-type and knockout females were bred to wild-type males for eight weeks. *Spag17* knockout females became pregnant and gave birth in their first pregnancy. However, they showed reduced litter size compared to wild-type females (Fig. 2A). Interestingly, one in five females died from obstructive labor, while the other four females became pregnant a second time, and three of them died during labor. Although one gave birth the second time, it did not become pregnant a third time (Fig. 2B).

**Figure 2:**
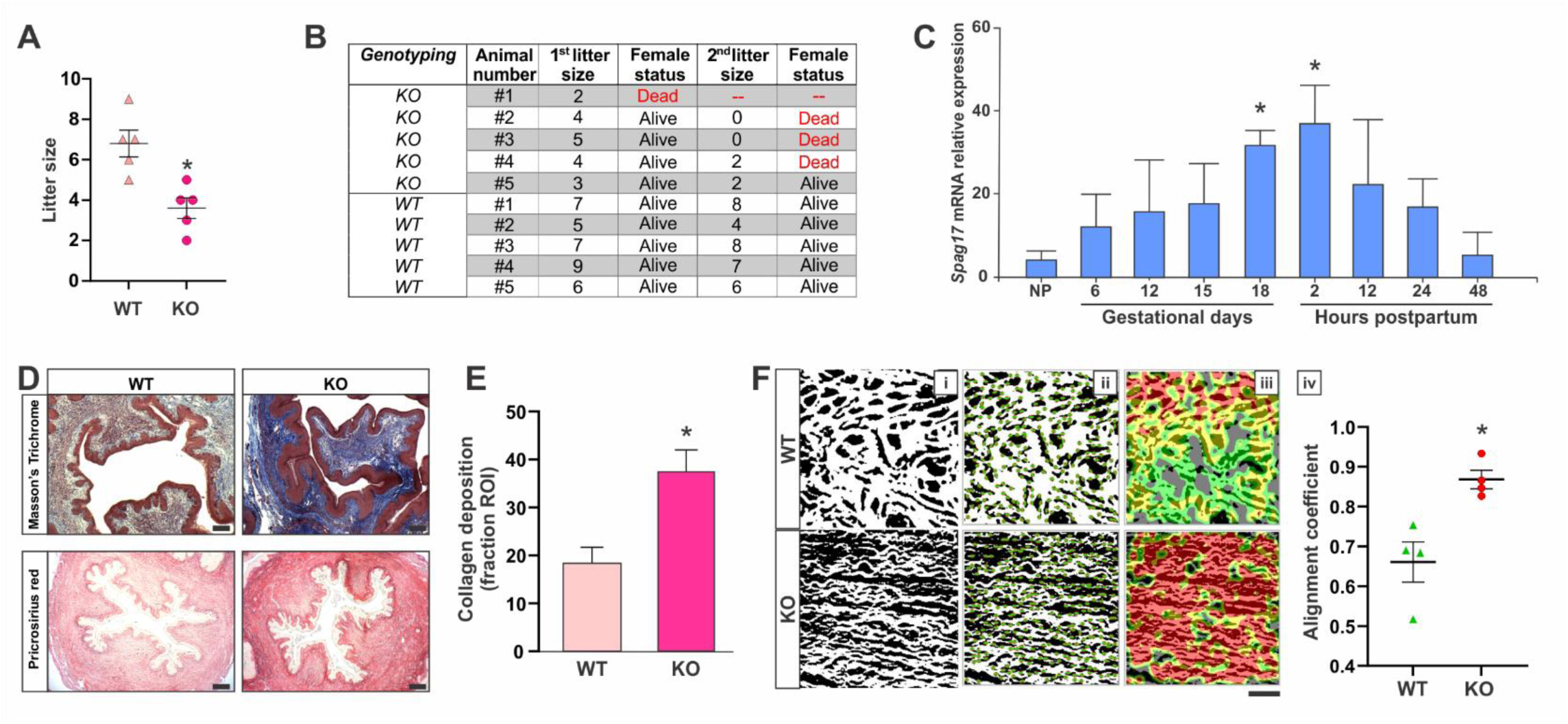
SPAG17 deficiency results in impaired fertility and heightened pregnancy complications. Young (6-weeks-old) females were mated with WT reproductive age males to evaluate fertility outcomes. (**A**) Quantification of litter size. Data are means ± SEM, n=5, *statistically differences vs. wild-type, p<0.05. (**B**) Quantification of litter size and female status during breeding protocol. *Spag17* knockout females have reduced litter size and severe parturition complications leading to maternal death. (**C**) *Spag17* relative expression in the mouse cervix at different gestational time points. Cervix tissue was collected from young wild-type female mice at different time points and analyzed for *Spag17* mRNA expression by qPCR. n=6 animals per time point. Data are means ± SEM. *Statistically differences vs. non-pregnant (NP), p<0.05. (**D**) Cervices from wild-type and *Spag17* knockout (n=6) mice were collected and processed for histological evaluation. Representative images showing Masson’s Trichrome and Picrosirius Red staining. Scale bars, 100 µm and 200 µm, respectively. (**E**) Quantification of Picrosirius Red staining between wild-type and *Spag17* knockout cervices. Data are means ± SEM, *p=0.02. (**F**) Quantification of collagen alignment. **i**, representative boundary mask analysis of the ROI area in PSR images; **ii,** representative images showing the angle measurement (with respect to the horizontal) using CFR mode, where a red dot indicates the center of fiber segments and a green line represents the fiber orientation at that point; **iii,** representative images displaying the heatmap, where red indicates well-aligned fibre regions; **iv,** quantification of the collagen alignment coefficient. Data are presented as means ± SEM, *p=0.01. Scale bar, 20 µm.

To further characterize this phenotype, we first evaluated the cervical expression of *Spag17* at different gestational times and in the postpartum period (2 to 48 h). An increase in *Spag17* mRNA expression during gestation was detected, reaching its highest level around 2 h postpartum, and declining between 12 and 48 h (n=6; Fig. 2C). During parturition, the cervix became more elastic and softer in preparation for labor. Immediately after labor, an extensive remodeling of the ECM takes place, allowing the cervix to recover [Barnum et al., 2017]. Remarkably, changes in *Spag17* expression seem to follow cervical homeostasis in preparation for labor, as the highest mRNA expression of *Spag17* was correlated with expected changes in the cervix such as dilation (widening of the cervix) and effacement (shortening of the cervix).

Next, we investigated whether changes in the ECM components such as collagen content were different in the knockout mice, which may explain the dystocia phenotype. We conducted comprehensive histological evaluations of the cervix in reproductive age female mice. Remarkably, both Masson’s trichrome staining and picrosirius red (PSR) staining revealed higher staining in the *Spag17* knockout tissues compared to the wild-type controls (n=6; Fig. 2D). Further quantitative analysis of the picrosirius staining demonstrated a marked increase in collagen deposition in the cervices of the knockout mice (n=5; Fig. 2E). Next, we investigated whether changes in collagen deposition were also accompanied with changes in collagen fiber alignment. Using CurveAlign (v. 5.0 beta) to analyze collagen fiber alignment in regions of interest, we found that knockout samples exhibit 1.3 times greater collagen fiber alignment compared to wild-type mice (Fig. 2F). These findings suggest that the dystocia phenotype may be attributed to defects in cervical dilation due to altered collagen composition and organization leading to increased matrix stiffness and fibrosis.

To further evaluate this possibility, we analyzed biomechanical properties of the cervix. Notably, we observed greater resistance in the tissue without breaking after applied forces (max load), along with increased maximum stress, stiffness, and modulus properties in *Spag17* knockout cervices compared to wild-type cervices (n=2; Fig. S3). These findings strongly support the notion that aberrant collagen composition and organization in the cervix lead to increased stiffness and fibrosis, which may explain the dystocia phenotype observed in the knockout females.

### *Spag17* knockout cervices show differential expression of genes associated with extracellular matrix remodeling, collagen, inflammation, aging and fibrosis

To further characterize the phenotype associated with cervical dysfunction, bulk RNAseq was conducted on cervical tissue from 6-to 10-week-old wild-type (n=4) and *Spag17* knockout (n=4) mice. Unbiased transcriptome profiling revealed differential expression of 793 genes in the knockout cervices. Of these DEGs, 398 were upregulated and 395 were downregulated (Fig. 3A). A heatmap correlation clearly demonstrated distinct clustering of DEGs between wild-type and knockout samples, highlighting the robust transcriptional differences (Fig. 3B). Gene set enrichment and gene ontology analysis show differences in genes associated with inflammation, aging, extracellular matrix, collagen, and fibrosis (Fig. S4A-C; Fig. S5A-C; Table S1-S3) terms from DEGs. Pathway analyses show enrichment of signaling pathways involved in the migration and chemotaxis of leukocytes, neutrophils, and granulocytes; cytokine and chemokine activity; receptor-ligand interactions; and components of the collagen-containing extracellular matrix and cytoskeletal organization (Fig. S6A-C). Moreover, Sankey diagrams illustrated a strong gene signature for specific pathways associated with fibrosis, inflammation, and aging/senescence mechanisms (Fig. 3C-E, respectively).

**Figure 3:**
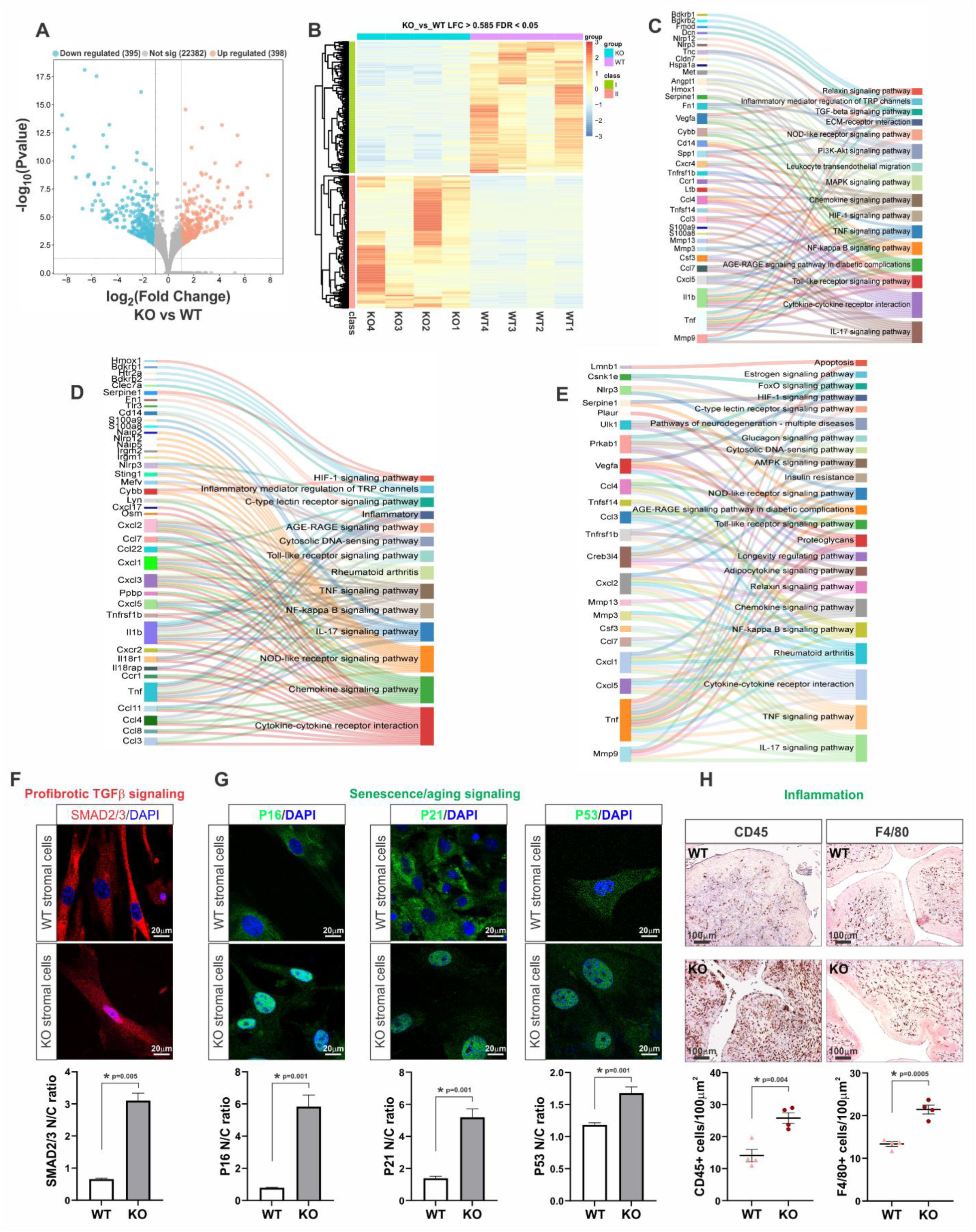
Spag17-deficiency drives a profibrotic, proinflammatory and aging/senescence-related signature in the cervix. Total RNA was extracted from WT and KO cervices (n=4) and subjected to RNAseq studies. (**A**) Volcano plot showing differential gene expression in *Spag17* KO cervices. (**B**) Heatmap of differentially expressed genes, red indicates overexpression, and blue indicates under-expression. Sankey plot of functional enrichment pathway analysis of (**C**) profibrotic, (**D**) proinflammatory, and (**E**) aging/senescence-associated signatures. Stromal cells from WT and KO cervices were cultured and used for immunofluorescence evaluation of profibrotic TGF-β signaling (**F**) and aging/senescence-associated markers (**G**). Data are means ± SEM, n=3, *p<0.05. Scale bar, 20 µm. (**H**) Immunolabeled detection of CD45 and F4/80 positive cells as inflammation markers in cervical tissues. Data are means ± SEM, n=4, *p<0.05. Scale bar, 100 µm.

To confirm these findings, stromal cells isolated from cervical tissue samples were cultured and analyzed using immunofluorescence to assess the activation of profibrotic signaling pathways, including TGF-β signaling, as well as markers of cellular senescence such as p21, p16, and p53. The results revealed a marked increase in the nuclear translocation of Smad2/3 proteins in knockout cells, indicating constitutive activation of this profibrotic signaling pathway (Fig. 3F). Moreover, increased nuclear translocation of the senescence markers p21, p16, and p53 in knockout cells was observed, consistent with a senescence signature (Fig. 3G).

To investigate the presence of cells involved in inflammation, immunohistochemistry was performed in wild-type and *Spag17* knockout cervix samples using anti-CD45 and anti-F4/80 antibodies. A notably high percentage of CD45 and F4/80 positive cells was observed in the knockout samples compared to wild-type (Fig. 3H), indicating heightened immune cell recruitment and inflammation in the knockout cervical tissue.

Collectively, these findings suggest that these molecular mechanisms drive the cervical phenotype observed in *Spag17* knockout mice and provide a framework for understanding the underlying biology of cervical dysfunction in this mouse model.

### Loss of SPAG17 leads to accelerated aging in the ovary

Single-cell RNAseq has been conducted by several groups to characterize ovarian aging [Isola et al., 2024, Jones et al., 2024, Wu et al., 2024; Zhou et al., 2024]. Changes in the molecular signatures of the different types of ovarian cells during physiological aging were defined. Remarkably, transcriptomic data from Isola and collaborators (2024) conducted in mouse tissues at single-cell resolution revealed a reduction in *Spag17* mRNA expression in aged ovaries, particularly in theca, endothelial and granulosa cells (Fig. 4A). Moreover, the expression of *Spag17* showed a negative association with *Col1a1* (Fig. 4B), *Col4a1* and *Acta2* (Fig. 4C), as well as other fibrosis-associated genes (Fig. S7). A list of proinflammatory genes also showed negative association with *Spag17* including *Ccl2*, *Ccl8*, *Lcn2*, *Il1b*, and *Il6* (Fig. 4D) and others (Fig. S8). In addition, specific senescence-related markers including *Cdkn2a* (p16), *Cdkn1a* (p21) and *Hmgb1* (p53) demonstrated an inverse association with *Spag17* expression in the aged ovaries (Fig. 4E and Fig. S9), suggesting that the physiological ovarian aging processes may be associated with lower expression of *Spag17*. Based on this observation, we predict that loss-of-function of *Spag17* promotes accelerated aging.

**Figure 4:**
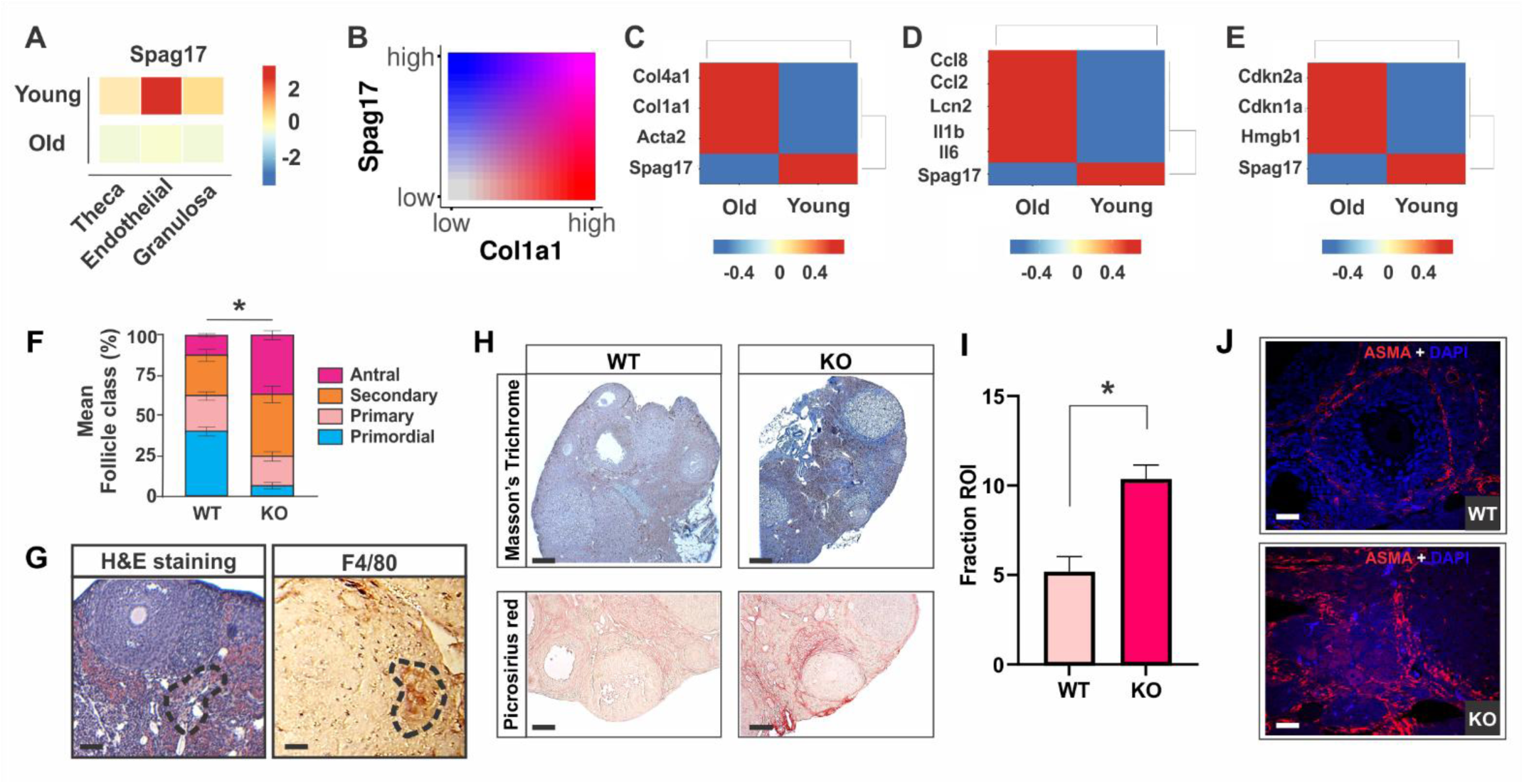
Age-related changes in Spag17 knockout ovaries. (**A**) Heatmap showing expression of *Spag17* in theca, endothelial and granulosa cells from data obtained from [Isola et al., 2024] comparing young and old mouse ovaries. (**B**) *Spag17* expression negatively correlates with *Col1a1* gene expression. Heatmap showing negative association of *Spag17* expression and genes linked to (**C**) profibrotic, (**D**), proinflammatory and (**E**) aging/senescence mechanisms in aged-ovaries. (**F**) Ovaries from young (5-week-old) WT and KO mice were collected and processed for histological H&E staining. Quantification of follicle classes detected as a percentage of the total ovarian follicle pool. Data are means ± SD, n=6, *p<0.05 (**G**) Representative images of H&E staining and F4/80 immunohistochemical-labeling (n=3). Dashed area indicates the presence of multinucleated macrophage giant cells. Scale bars, 100 µm. (**H**) Representative images of Masson’s Trichrome and Picrosirius Red-stained ovarian sections show increased collagen deposition in *Spag17* knockout ovaries. Scale bars, 200 µm. (**I**) Quantification of Picrosirius Red-staining between WT and KO ovaries. Data are means ± SEM, n=6, *p<0.05. (**J**) Representative images of immunofluorescent detection of ASMA (red). DAPI (blue) was used for nuclear labeling, n=3.

To examine this possibility, we next conducted a comprehensive evaluation of the ovarian phenotype in *Spag17* knockout mice. Advanced female age is associated with decreased oocyte quality and quantity, fibrosis, and the accumulation of a distinct population of multinucleated macrophage giant cells (MNGCs), which are linked to chronic inflammation [Duncan et al., 2018; Briley et al., 2016; Umehara et al., 2018]. Therefore, we began our analysis with a histological evaluation of the ovaries. Notably, *Spag17* knockout ovaries exhibited architectural differences compared to wild-type ovaries of the same age. Specifically, a reduced number of primordial follicles was observed (n=5; Fig. 4F). This finding suggests that fewer oocytes are available for fertilization in knockout ovaries compared to wild-type controls, potentially contributing to the reduced litter sizes observed in knockout mice (Fig. 2A).

Moreover, *Spag17* knockout ovaries displayed the presence of MNGCs, a hallmark of chronic inflammation. This was evident from H&E staining and immunohistochemistry using anti-F4/80 antibody (n=6; Fig. 4G). In addition, Masson’s trichrome and picrosirius red staining revealed significantly increased collagen deposition in the stroma (Fig. 4H-I). This phenotype was accompanied by elevated expression of the myofibroblast marker α-smooth actin (ASMA), detected via immunofluorescence (n=3; Fig. 4J). To further refine the characterization of this fibrotic phenotype, PSR-stained ovarian samples (n=5) were analyzed using the FibroNest™ digital pathology platform, which employs quantitative single-fiber artificial intelligence (AI) image analysis [Kostadinova et al., 2024]. This novel methodology provides an extensive, comprehensive and precise assessment of the intricate changes in collagen content, fiber morphometry, and architecture (Fig. 5A). The resulting fibrotic phenotypic heatmap revealed a consistent increase in the severity of all quantitative fibrosis traits (qFT) in *Spag17* knockout ovaries compared to wild-type controls (Fig. 5B and Fig. S10). Additionally, a higher collagen composite score was observed (Fig. 5C), indicating increased collagen deposition. Notably, single-fiber analysis highlighted the enrichment of densely packed fibers forming complex structures in the stromal region of the ovaries (Fig. 5A; Fig. S11). The quantification of fine and assembled fibers demonstrated a higher complex fiber structure in the knockout ovaries (Fig. 5A and D-F). Moreover, texture analysis revealed a more intricate collagen architecture (Fig. 5A and G). Consistent with these findings, the consolidation of all 195 principal phenotypic traits resulted in a higher fibrosis composite score in these samples (Fig. 5H). Together, these findings suggest that loss-of-function of *Spag17* promotes accelerated aging and fibrosis in the ovaries. Therefore, SPAG17 may be an important modulator of aging-related mechanisms.

**Figure 5:**
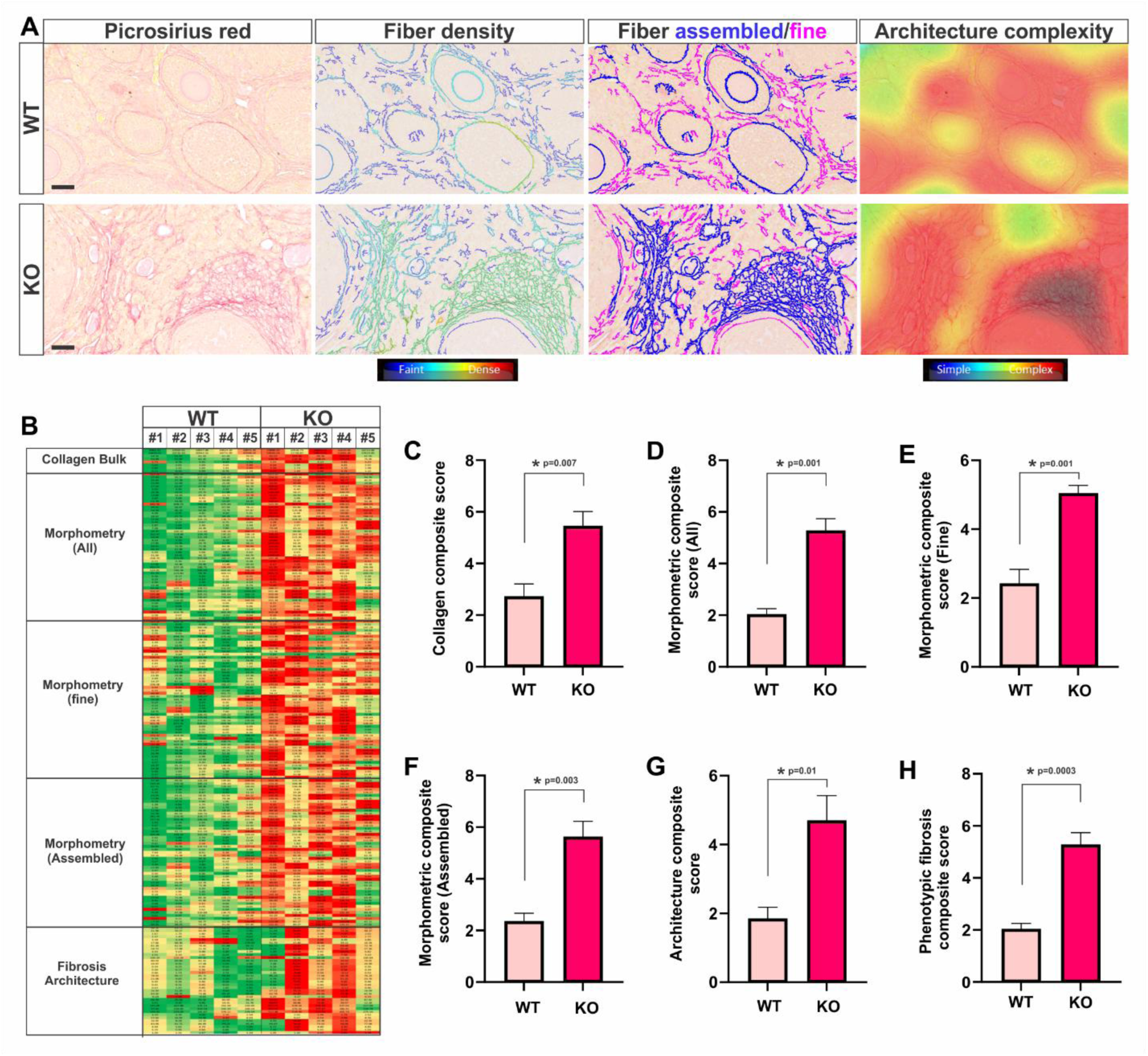
Loss of SPAG17 leads to fibrosis in the ovarian tissue. PSR-stained ovarian tissues from WT and KO mice were analyzed using the FibroNest™ digital pathology platform (n=5). (**A**) Representative images showing characterization of collagen deposition and fiber morphology and architecture. Scale bars, 50 µm. (**B**) Heatmap showing the extent of fibrosis severity, ranging from green (faint and fine collagen) to red (dense and complex collagen). Each column represents a sample, and each row corresponds to a different quantitative fibrosis trait (qFT) contributing to the collagen, morphometric, and architectural sub-phenotypes. Quantification of composite score based on the major qFTs of each sample group: (**C**) Collagen composite score; (**D**) Morphometric composite score (ALL); (**E**) Morphometric composite score (Fine); (**F**) Morphometric composite score (Assembled); (**G**) Architecture composite score; (**H**) Phenotypic fibrosis composite score calculated using a total of 195 principal phenotypic traits. Data are means ± SEM, *p<0.05.

## DISCUSSION

Morphological changes in female reproductive tissues, such as inflammation and fibrosis, are increasingly recognized as early hallmarks of reproductive aging [Balough et al., 2024]. The cellular and molecular mechanisms driving these changes are complex and remain only partially understood. Emerging evidence points to a multifactorial process involving various cell types, organelles, and signaling pathways [Duncan et al., 2018].

Our study reveals that loss-of-function of *Spag17* promotes constitutive activation of signaling pathways linked to immune responses, fibrosis, and senescence, leading to immune cell infiltration, increased collagen deposition and fibrosis. These morphological and molecular changes suggest *Spag17* is a key regulator of molecular mechanisms underlying reproductive aging.

The occurrence of cellular senescence has been tightly associated with accelerated aging. Senescence can be triggered by several signaling pathways including kinases MAPKK3/ MAPKK6, oncogenic signaling or loss of tumor suppressors (RAS, MYC, PI3K) and TGF-β, which activates common regulators including p53, p16 and p21 [Muñoz-Espín and Serrano, 2014]. Cellular senescence increases the secretion of proinflammatory cytokines from the SASP [Tchkonia et al., 2013], impacting tissue homeostasis and contributing to a profibrotic response [Selman & Pardo, 2021].

Here, we show that the absence of SPAG17 in the cervix results in significant enrichment of gene ontologies related to leukocyte, neutrophil, and granulocyte migration and chemotaxis; cytokine and chemokine activity; receptor-ligand interactions; collagen-containing extracellular matrix; metalloendopeptidase activity; and cytoskeletal structure among others. Furthermore, key signaling pathways, including the TGF-β pathway, were constitutively activated, along with downstream effectors such as p53, p21 and p16. Consequently, *Spag17* knockout females exhibit increased fibrosis in the cervix, leading to complications during parturition and maternal death—paralleling pregnancy complications observed in aged women [Laopaiboon et al., 2014]. Similarly, a robust aging phenotype was detected in the ovaries of *Spag17* knockout mice that mimic the phenotypes occurring during physiological ovarian aging [Zhang et al., 2020; Wu et al., 2023; Isola et al., 2024, Jones et al., 2024; Zhou et al., 2024].

The cervix and ovaries are composed of a heterogeneous mix of various cell types that differ in gene expression and function. A recent study demonstrated that fibroblasts play central—and highly organ-specific—roles in remodeling the female reproductive tract. They orchestrate extracellular matrix reorganization and inflammation throughout the reproductive lifespan, driving the gradual, age-related development of fibrosis and chronic inflammation. This process seems to be mediated through their cell-cell communication network via ligand-receptor interactions [Winkler et al., 2024]. A limitation of our study is the absence of a detailed analysis examining the specific roles of individual cell types within the cervix and ovaries that contribute to the accelerated aging phenotype. Given that SPAG17 is expressed in multiple cell types in these tissues, future research will aim to address this limitation by exploring the unique functions and contributions of each cell type to the observed phenotype.

Collectively, our results suggest that SPAG17 plays a pivotal role in mitigating the effects of aging in the female reproductive tract by regulating ECM remodeling, inflammation, and fibrosis. The loss of SPAG17 disrupts these processes, leading to structural and functional impairments that manifest as accelerated aging, reduced fertility and obstructive labor (Fig. 6). Given the rising average maternal age worldwide and the associated increase in reproductive complications, understanding the molecular mechanisms underlying reproductive aging is crucial. SPAG17 may represent a novel therapeutic target for mitigating aging-related dysfunction, potentially improving reproductive outcomes in women of advanced maternal age.

**Figure 6:**
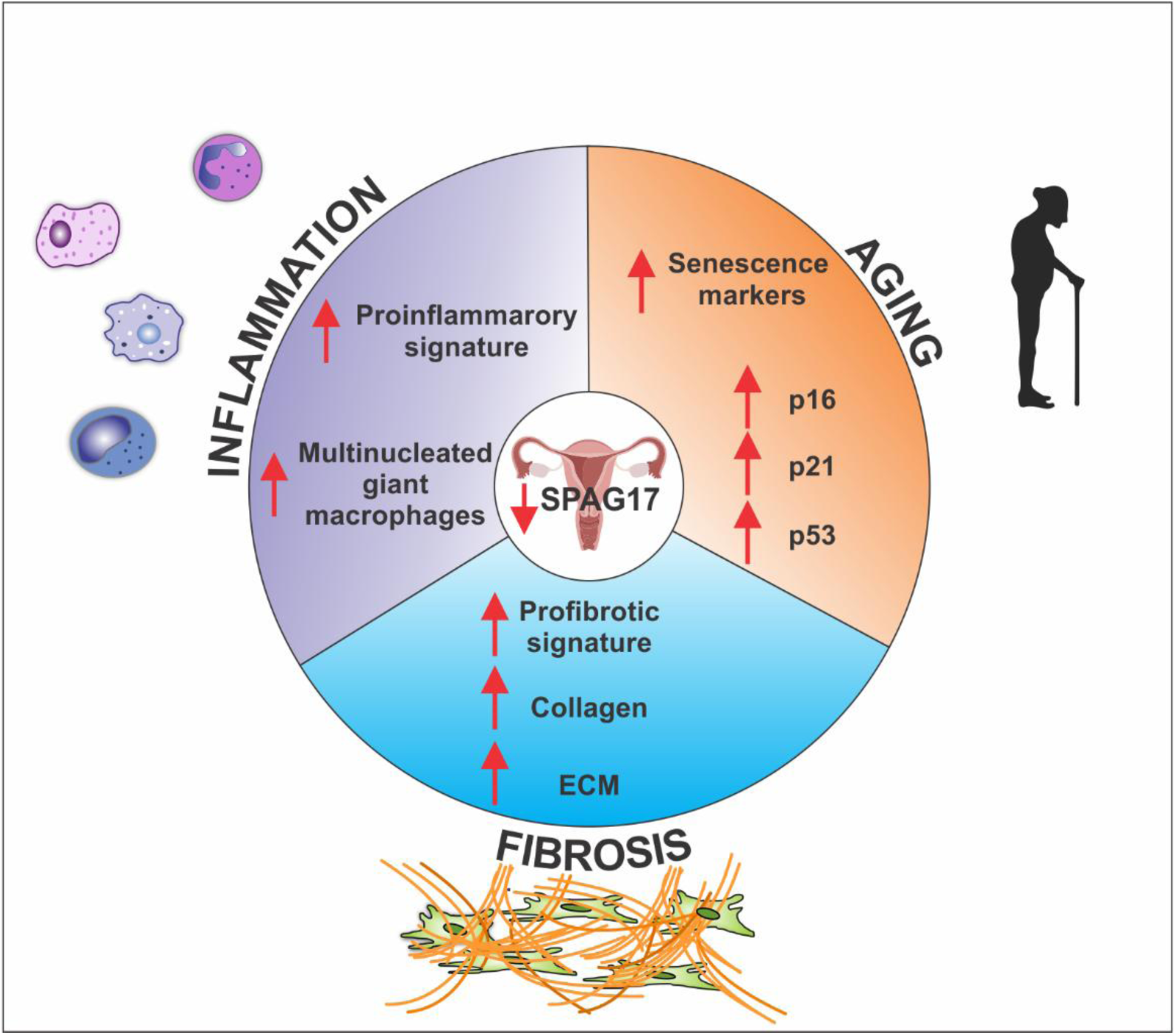
Regulation of Female Reproductive Aging Through the Spag17 Gene. Schematic representation of the mechanisms modulated by *Spag17* that lead to an aging phenotype. Reduced expression or loss of SPAG17 function promotes inflammation, ECM remodeling, fibrosis, and senescence, resulting in accelerated aging, ultimately leading to reduced fertility and obstructed labor.

## MATERIALS AND METHODS

### In silico analysis of publicly available RNAseq data

Single-cell RNAseq datasets from human atlas; Guo et al., 2022 and Isola et al., 2024 were used for in silico analysis to determine the expression of *SPAG17* in the female reproductive tract. To perform this analysis, we first used the Fenn server from VCU to call for the SRR list from the samples deposited in GEO by the authors. Next, the samples were processed using Cell Ranger v7.2 and visualized and analyzed by Seurat v5 program. The previously published dataset [Isola et al., 2024] was obtained and downloaded from a Shiny-based web app (https://omrf.shinyapps.io/ovarianagingscatlas/). Data were analyzed using bubble plots and heatmaps with gene expression patterns from young and old ovaries. Shinycell R package, graphics were set using the R/Bioconductor packages clusterProfiler [Yu et al. 2012] and Pathview [Luo and Brouwer, 2013].

Other data used for the analyses described in this manuscript were obtained from the GTEx Portal on 07/07/24.

### Animals

All animal studies were conducted in accordance with protocol AM10297 approved by the Virginia Commonwealth University Institutional Animal Care and Use Committee. Mice were housed in a controlled barrier facility at VCU under constant temperature, humidity, and light (14 h light/10 h dark). Mice were provided with food and water ad libitum. Females from 5-weeks to 5-months old were used in this study. *Spag17*-mutant mice were generated by crossing heterozygous *B6N(Cg)-Spag17 tm1b(KOMP)Wts1/J* (Stock No. 026485) mice from Jackson Laboratories. The resulting homozygous offspring (*Spag17/Sox2-Cre* knockout) mice carry a deletion in exon 5 that leads to a premature stop codon and absent *SPAG17* expression [Kazarian *et al*., 2018; Sapao et al., 2023]. The wild-type mice used as controls share the same genetic background as the *Spag17/Sox2-Cre* knockout mice. To ensure consistency and minimize genetic variation, we typically use mice from the same litter or mice derived from the same breeding line. The total number of animals used for these studies were wild-type n=47 and knockout n=59.

### Mouse ovary and cervix harvest

Wild-type and *Spag17* knockout female mice were euthanized according to protocol AM10297. Mice were then prepped on a harvesting pad and fur was cleaned thoroughly with 70% alcohol. Incisions were made to the abdominal region, after which ovaries, uterine horns, cervix, and vagina were removed. Ovaries were separated, and one was fresh frozen in dry ice and stored at -80° C and the other was fixed in 10% formalin for histological studies. Cervices were either fresh frozen or fixed.

### β-Galactosidase staining

To visualize the expression of the *Spag17* gene, ovarian and cervical tissues were collected from heterozygous *Spag17/Sox2-Cre* females (n=3) and stained using a beta-Gal Reporter Gene Staining kit (Sigma, Saint Louis, MO). After tissues were removed from the mice, they were washed with PBS to remove blood. Then tissues were fixed in 1X fixation buffer with 20% formaldehyde and 2% glutaraldehyde in PBS for 1 h. After fixation, they were quickly rinsed with PBS and then incubated in PBS for 30 min. Ovarian and cervical tissues were then placed in the staining solution with 200 mM MgCl2, 400 mM potassium ferricyanide, 400 mM potassium ferrocyanide, 20 mg/mL of X-gal, and 0.02% Triton X-100 in 1X PBS overnight in the dark at 37°C. To avoid endogenous β-Gal activity, pH was kept around 7.5 to 8.5. The stained tissues were rinsed with PBS for 30 min and fixed in 4% PFA overnight at 4°C. Following the fixation and PBS rinsing, tissues were sectioned at a thickness of 40 µm in 0.1M phosphate buffer with a vibratome (Leica, Banockburn, IL). Images were acquired using a Discovery V20 Stereo zoom fluorescence stereoscope (Zeiss).

### Immunohistochemistry

The tissue-sectioned slides (WT n=4-6 and KO, n=4-6) were subjected to deparaffinization using 100% xylene, followed by rehydration through a series of decreasing ethanol concentrations. Endogenous peroxidase was removed by treating the slides with 3% H_2_O_2_ in methanol for 30 min. Antigen retrieval was conducted by immersing the slides in citrate buffer at 95°C for 20 min. After multiple washes with PBS, the slides were incubated in a blocking solution comprising 10% goat serum and 0.02% triton X-100 in 1X PBS for 1 h at room temperature. An overnight incubation at 4°C was performed using a primary anti-SPAG17 rabbit polyclonal antibody [Teves et al., 2013; Teves et al., 2015] diluted at 1:200, anti-F4/80 rat monoclonal antibody (1/50 dilution, ThermoFisher #14-4801-82), or anti-CD45 rat monoclonal antibody (1/100 dilution, ThermoFisher #14-0451-82). Subsequently, the slides were exposed to a biotinylated secondary anti-rabbit or anti-rat antibody (Vector Laboratories, Burlingame, CA, USA) for 1 h at room temperature. Following this, the slides were washed with PBS and treated with Vectastain Avidin-Biotin-Complex (ABC) reagent (Vector Laboratories Inc., Burlingame, CA, USA) for 30 min at room temperature. After several PBS washes, staining was developed using ImmPACT Diaminobenzidine (DAB) for 2 min and the reaction was stopped by rinsing with tap water. Wild-type and *Spag17* knockout samples were simultaneously processed to ensure consistent conditions. Lastly, the stained slides were dehydrated through an ethanol and xylene gradient and mounted using Vectastain (Vector Laboratories Inc., Burlingame, CA, USA) mounting media. Image analysis was performed using a digital camera (Olympus QColor5), attached to an Olympus BH-2 microscope (Olympus, Center Valley, PA).

### Fertility assessment

To assess fertility, young (6 weeks old) virgin females (WT n=5 and KO, n=5) were individually housed with 3-month-old WT males of proven fertility. Breeding cages were checked daily for plugs, and the number of litters, number of pups born, and pup and dam survival were recorded.

### Mouse cervix collection and Spag17 mRNA expression at different gestational time points

Tissue collection was conducted at the University of Texas Southwestern Medical Center (Dallas, TX). Research protocols using mice were reviewed and approved by the Institutional Animal Care and Use Committee at the University of Texas Southwestern (UTSW) Medical Center in Dallas, Texas. The animal procedures were performed under the standards of humane animal care following the NIH Guide for the Care and Use of Laboratory Animals. C57B6/129sv wild-type nulliparous mice were housed under a 12 h-light/12 h-dark cycle at 22 °C. Pregnancy was confirmed with the presence of a vaginal plug (considered day 0), with birth typically occurring early morning on day 19. Cervical tissues were collected from non-pregnant (NP) mice, as well as on gestation days 6 (d6), 12 (d12), 15 (d15), and 18 (d18). Additionally, postpartum cervical tissues were collected at 2, 12, 24, and 48 h after birth (n=6 per gestation group). The collected tissues were immediately frozen. After tissue collection, standard procedures were followed for total RNA extraction and RT-PCR. The *Spag17* mRNA expression was assessed using the following primer sets: forward primer (5’-CTTGAAGTGTCAACTTCTCC-3’, located within exon 4) and reverse primer (5’-CCAAGCTCAGTCATAATGGCC-3’, in exon 6). Quantitative PCR reactions were run in 96-well plates with an end volume of 20 µl per sample, containing 10 µl SiTaq Universal SYBR green supermix (BIO-RAD, Hercules, CA, USA), 300 nM of each primer, 50 ng/ml of cDNA, and a realplex mastercycler (Eppendorf, Hamburg, Germany). The thermocycler program included an initial denaturation of 95 °C for 2 min, followed by 40 cycles of 95 °C for 15 sec, and an annealing and elongation stage of 62 °C for 1 min. Melt curve analysis was performed at the end of each run to check for multiple peaks indicative of non-specific amplification. Cycle threshold data (CT) were normalized relative to 18SrRNA for each plate (ΔCT).

### RNA Sample Collection and Preparation

Total RNA was isolated from wild-type and *Spag17* knockout mouse cervices using TRIzol® (Life Technologies, Inc., Grand Island, N.Y.) as previously reported [Teves et al, 2015]; 1 ml of RNAlater was used to preserved RNA integrity until isolation. From 100 ng of input RNA, libraries for transcripts abundance and sequencing were performed using the NEBNext® UltraTM RNA Library Prep Kit (Illumina, San Diego, CA, USA) following the manufacturer’s instructions on a NovaSeq 6000 instrument.

### RNAseq analyses in wild-type and Spag17 knockout cervix samples

The RNAseq analysis was performed for 8 samples (4 KO and 4 WT) using the OneStopRNAseq package [Li et al., 2020]. Specifically, FastQC [Andrews, 2010] and MultiQC [Ewels et al., 2016] were used for raw reads quality control and QoRTs [Hartley & Mullikin, 2015] for post-alignment quality control. In this regard, reads were aligned to the reference genome assembly mm10 with star_2.7.5a [Dobin et al., 2013] and annotated with gencode.vM25.primary_assembly [Harrow et al., 2012]. Aligned exon reads were counted toward gene expression with featureCounts_2.0.0 [Liao et al., 2014] with default settings except for paired-end strict-mode analysis parameters (e.g. ‘-Q 20 --minOverlap 1 --fracOverlap 0 -p -B-C’). Differential expression (DE) analysis was performed with DESeq2_1.28.1 [Love et al., 2014]. Within DE analysis, ‘ashr’ was used to create log2 Fold Change (LFC) shrinkage for each comparison, which is used to avoid extreme LFC values [Stephens, 2016]. Significantly differentially expressed genes (DEGs) were filtered with the criteria FDR < 0.05 and absolute log2 fold change (|LFC|shrunken) > 0.585. Heatmaps were created with pheatmap [Kolde 2012.]. Gene set enrichment analyses were performed with GSEA [Subramanian et al., 2005] and gene ontology/pathway graphics were set using the R/Bioconductor packages clusterProfiler [Yu et al. 2012] and Pathview [Luo and Brouwer, 2013]. Alternative splicing analysis was performed with rMATS_4.1.0 [Shen et al., 2014].

### Histological evaluation

Fixed tissues (WT n=6 and KO, n=6) were paraffin-embedded and sectioned into 5 µm slices. Hematoxylin and eosin staining was performed using standard procedures and captured with a digital camera (Olympus QColor5), attached to an Olympus BH-2 microscope (Olympus, Center Valley, PA).

### Picrosirius Red and Masson’s Trichrome staining

For Picrosirius Red (PSR) staining, tissue sections (WT n=6 and KO, n=6) were deparaffinized in two 100% xylene solutions for 5 min each. The slides were then rehydrated in a series of ethanol baths (2 times 100%, 70%, 35%) for 5 min each, and lastly washed in a bath of PBS. The slides were then immersed in a PSR solution consisting of Sirius Red (Direct Red 80, Abcam) in an aqueous saturated solution of picric acid (Abcam 1.2%) at 0.1% w/v for 60 min at room temperature. Slides were then rinsed for 20 sec (two times) in 5% acetic acid. Lastly, slides were dehydrated in two changes of 100% alcohol for 2 min each and mounted with VECTA mount (Vector laboratories).

Masson’s Trichrome (MTC) staining kit (Polysciences, Inc., Washington, PA) was used following the manufacturer’s instructions. Briefly, tissue sections (WT n=6 and KO, n=6) were deparaffinized and rehydrated as mentioned above. Slides were submerged in Bouin’s fixative and placed on a slide warmer to incubate for 60 min at 60°C. These slides were then washed in running tap water to remove the yellow picric acid for 5 min. Weigert’s iron hematoxylin working solution (a combination of Weigert’s Hematoxylin A and Weigert’s Hematoxylin B at a 1:1 ratio) was placed on the slides for 1 min and washed under running tap water for 5 min and transferred to 1 min rinse in distilled water. Next, the slides were stained in Beibrich Scarlet-Acid Fushsin for 30 sec, followed by a 1 min rinse in distilled water. The rinsed slides were incubated with phosphotungstic/phosphomolybdic acid for 10 min and then incubated in Aniline Blue for 30 min. After rinsing the slides in distilled water, they were incubated in 1% acetic acid for 30 sec and rinsed. Lastly, the slides were dehydrated in ethanol baths (95%, 100%) for 2 min each and submerged in xylene before being mounted with VECTA mount (Vector Laboratories, Inc., Burlingame, CA, USA).

PSR-stained slides were imaged using the Olympus BH-2 microscope and Olympus (Q-COLOR 5 RTV) camera with cellSens Dimension 1.15 software. The entire tissue was imaged in composite sections under 4 and 10x magnification. Regions of interest (ROI) were selected within each partial image and analyzed using the cellSens Dimension 1.15 software count and measure tool. First, we selected an intensity threshold that was specific to the intensity of the staining. Applying these features to imaged samples, we calculated the Area Fraction ROI (%). This value measures the area within the ROI that meets the intensity criteria. These values were collected for the various ROIs selected for each section. Next, the differences in % ROI between wild-type and *Spag17* knockout samples were evaluated.

### High Resolution Digital Pathology Quantitative Image Analysis

Picrosirius red-stained tissue sections (WT n=5 and KO n=5) were imaged at 40x magnification in brightfield using a Vectra Polaris™ Automated Quantitative Pathology Imaging System. Images were then analyzed with the FibroNest™ digital pathology platform (PharmaNest, Princeton, NJ, USA), which employs single-fiber quantitative image analysis and artificial intelligence (AI) for automated phenotypic quantification of fibrosis. This method enables highly sensitive quantitative fibrosis analysis by quantifying and calculating various quantifiable traits in collagen content, morphology, and fibrosis architecture to establish algorithm-based quantitative fibrosis scores. These scores have been validated against different fibrosis scoring methods [Soon & We, 2021; Briand et al., 2021; Nakamura et al., 2022; Inia et al., 2023; Wang et al., 2023]. After pre-processing steps including color normalization, standardization, and segmentation to eliminate staining variability, scanning artifacts, compression artifacts, and other sources of noise, each collagen fiber was identified and segmented as an individual object using a combination of specialized thresholding and AI methods. To account for the evolution of collagen fibers from faint and simple structures to more complex and networked ones, fibers were classified into fine and assembled fibers based on the number of skeleton nodes in their individual skeletons. Assembled collagen fibers are characterized by a complex skeleton with a high number of nodes and branches (>50 branches). Fine collagen fibers are less complex (<50 branches) and typically at the origin of fibrosis. Texture analysis was applied on the segmented collagen image layer to generate additional architectural quantitative parameters. Next, continuous severity scores for collagen deposition content (8 traits), fiber morphometry (151 traits), and organization of the fibers and architecture (36 traits) were calculated. Finally, a total of 195 principal phenotypic traits were combined to establish a normalized phenotypic fibrosis composite score (Ph-FCS). This approach allowed for the simultaneous assessment of architectural and morphological changes as well as quantity of collagen, providing a more comprehensive assessment of fibrosis in the analyzed tissues.

### Collagen alignment determination in cervical tissues

The orientation and alignment of collagen fibers were measured using CurveAlign v. 5.0 beta (Laboratory for Optical and Computational Instrumentation, University of Wisconsin, Madison, Wisconsin) following protocol from Liu et al., 2017 [Bredfeldt et al., 2014; Liu et al., 2017] on PSR-stained wild-type and *Spag17* knockout cervical tissue (n=4). Images were processed using ImageJ, undergoing background subtraction and binary conversion for optimal analysis. The ROI manager feature of the CurveAlign program was employed to compare collagen fiber alignment in the cervical tissue. Three, 256×256 pixel ROIs areas per sample were analyzed. CurveAlign with “Cropped Rectangular ROI” analysis was performed with a coefficient of 0.06. Fiber alignment was reported as values from 0 to 1, with 1 indicating perfectly aligned fibers, and smaller values representing more randomly distributed fibers. An unpaired t-test was performed to compare the mean of collagen fiber alignment between the samples.

### Biomechanic studies in cervical tissue

Wild-type (n=2) and *Spag17* knockout (n=2) females were used for biomechanical studies following protocol from Barnum and collaborators [Barnum et al., 2017]. Briefly, the cervical tissue was collected and laid flat to expose the lumen. The ends were affixed between two pieces of sandpaper for gripping, such that a uniaxial tensile load on the grips would simulate dilation of the cervical canal (loading occurred perpendicular to the proximal-distal direction). The tissue was continually immersed in PBS till the start of mechanical testing. A custom laser device was used to measure the cross-sectional area at a minimum of two locations. The approximate variability of tissue cross-section was within 20% of the average area. The cervix was then placed in custom fixtures to grip it at both ends. The cervix was then tested in tension using an Instron 5848 testing system (Instron Corp., Norwood, MA) using a standard protocol consisting of a preload of 0.005 N followed by a hold of 5 min and then a ramp to failure at mm/min. The location of failure was recorded for each sample.

### Stromal cell collection and culture

Stromal cells were isolated from the cervix of wild-type (n=3) and knockout (n=3) mice following a protocol adapted from Inada et al. (2006) [Inada et al., 2006]. Briefly, cervical tissue was removed from the animals, slit longitudinally, and placed in conical tubes containing 3 mL of 0.5% trypsin (Life Technologies Corporation, Grand Island, NY) and 2.5 mg/mL DNase I (STEMCELL Technologies, Vancouver, BC, Canada) for one hour at 37°C. The tubes were vortexed every 10 minutes to aid tissue digestion. After trypsinization, the tubes were vortexed for 10 sec. The supernatant containing epithelial cells was discarded, and the pellet was washed twice with 5 mL of Hanks’ Balanced Salt Solution (HBSS) (Life Technologies Corporation, Grand Island, NY) to remove any remaining epithelial cells. The remaining tissue was then transferred to a petri dish, cut into smaller pieces with a blade, and transferred back to a conical tube for an additional 30–45-min treatment with 0.5% trypsin and 2.5 mg/mL DNase I at 37°C. After this second trypsinization, the tubes were vortexed for another 10 sec, and the supernatant was transferred to a new tube and centrifuged for 5 min at 2000 rpm. The supernatant was then removed, and the pellet containing stromal cells was resuspended in 4 mL of culture media containing DMEM-F12 (ATCC, Manassas, VA), 10% FBS (Life Technologies Corporation, Grand Island, NY), and 1% penicillin-streptomycin. Cells were transferred to collagen-coated 8-well chamber slides (Falcon, Corning Inc., NY) and incubated at 37°C in a humidified atmosphere of 5% CO₂ in air. The culture media was changed every other day until the cells reached confluence, at which point they were used for immunofluorescence studies.

### Immunofluorescence

8-well chamber slides containing stromal cells from wild-type (n=3) and knockout (n=3) were fixed in 4% formalin (Sigma Aldrich, St. Louis, MO) and processed for immunolabelling using anti-Smad2/3 (1/100 dilution, Cell Signaling #5678), anti-p53 (1/200 dilution, Cell Signaling #2524), anti-p15/16 (1/50 dilution, Santa Cruz #sc-377412) and anti-p21 (1/50 dilution, Santa Cruz #sc-6246) antibodies. After washing, cells were incubated with anti-mouse Alexa Fluor 488-labeled (1/3000) or anti-rabbit Cy3-labeled (1/5000), secondary antibodies (Jackson ImmunoResearch Laboratory Inc., Grove, PA) for 1 h at room temperature, washed with PBS and mounted with VectaMount with DAPI (Vector Laboratories, Inc., Burlingame, CA, USA), and sealed with nail polish. Images were captured using a Zeiss LSM 700 confocal laser-scanning microscope. Images were analyzed using Zen10 and ImageJ software. The nuclear/cytoplasmic ratios for Smad2/3, p16, p21 and p53 were determined using ImageJ software. Total cell area was established from bright field images. Immunofluorescence images with blue (nucleus) and red or green channels depending on the secondary antibody used were separated. The blue channel determined the total nuclear area, while the red or green channels were used to quantify fluorescence intensity for the mentioned proteins. Fluorescence in the cytoplasmic region was derived by subtracting nuclear fluorescence from the fluorescence in the entire cell, then divided by the respective area to normalize. Subsequently, the nucleus’s fluorescence intensity was divided by cytoplasmic fluorescence intensity to calculate the N/C ratio.

Slides containing the sectioned tissue (WT n=3 and KO, n=3) were deparaffinized in 100% xylene and then rehydrated by decreasing the concentration of the ethanol. Antigen retrieval was performed in citrate buffer at 95 °C for 20 min. After washing several times in PBS, slides were incubated with 1% Triton X-100 (Sigma-Aldrich, St. Louis, MO, USA) for 5 min at 37 °C, washed with PBS three times, and blocked at room temperature for 1 h with 10% goat serum (Vector Laboratories, Inc., Burlingame, CA, USA) in PBS. Then, incubated overnight with anti-ASMA (Abcam # ab5694) antibody at 4 °C. After washing, incubated with anti-rabbit Cy3-labeled secondary antibody (Jackson ImmunoResearch Laboratory Inc., Grove, PA, USA) for 1 h at room temperature. Slides were subsequently washed with PBS, mounted with VectaMount with DAPI (Vector Laboratories, Inc., Burlingame, CA, USA), and sealed with nail polish. Images were captured using a Zeiss LSM 700 confocal laser-scanning microscope. Images were analyzed using Zen10 and ImageJ software.

### Follicle classification and counting

Follicle classification and counting were performed in wild-type and *Spag17* knockout ovaries. Serial sections from 5-weeks-old WT (n=6) and *Spag17* KO (n=6) were stained with Hematoxylin and Eosin. Then, sections were imaged in brightfield using a digital camera (Olympus QColor5), CellSens imaging software (Olympus), and an Olympus BH-2 microscope (Olympus, Center Valley, PA). Then follicles were classified and counted as previously described [Briley et al., 2016]. Primordial follicles were classified as oocytes surrounded by an incomplete or complete layer of squamous granulosa cells. Primary follicles were characterized as oocytes surrounded by a single layer of cuboidal granulosa cells. Primordial and primary follicles were counted irrespective of whether the oocyte nucleus was visible in the section. Secondary follicles were classified as oocytes surrounded by more than one layer of cuboidal granulosa cells. Antral follicles were classified by the presence of a visible antrum and a cumulus-oocyte complex surrounded by multiple layers of mural cells. Only secondary or antral follicles with an oocyte nucleus present in the section were included to avoid duplicate counting of these large later-stage follicles.

### Statistical analysis

Statistical analysis was conducted using Prism10 software by GraphPad (San Diego, CA, USA). For the comparison of two data groups, a t-test was employed, while a one-way ANOVA with Bonferroni corrections was utilized for the comparison of multiple data groups. The data were presented as means ± standard errors. The normality of the data was assessed using the D’Agostino and Pearson omnibus normality test. The significance between samples was determined at a p-value < 0.05.

Datasets were compared using analysis of variance (with Tukey’s or Dunnett’s multiple comparisons posttest) or otherwise when specified.

## Funding

CONAHCYT for a sabbatical stay at Teve’s Lab (CC-F).

R01HD105752 (FED)

R01 NS128104 (AL) R21 NS126611 (AL)

MCV Foundation Blick Scholarship (AL)

## AUTHOR CONTRIBUTIONS

Conceptualization: MET

Methodology: VE, MD, PS, LMTN, MEG, SC, CC-F, AL, LC, MP, FED

Investigation: SH, MM, FED, MET

Visualization: MET

Supervision: FED, MET

Writing—original draft: VE, MD

Writing—review & editing: JFS, JV, FED, MET

Competing interests

Authors declare that they have no competing interests.

Data and materials availability

All data are available in the main text or the supplementary materials.

## Supplementary Materials

**Figure S1:**
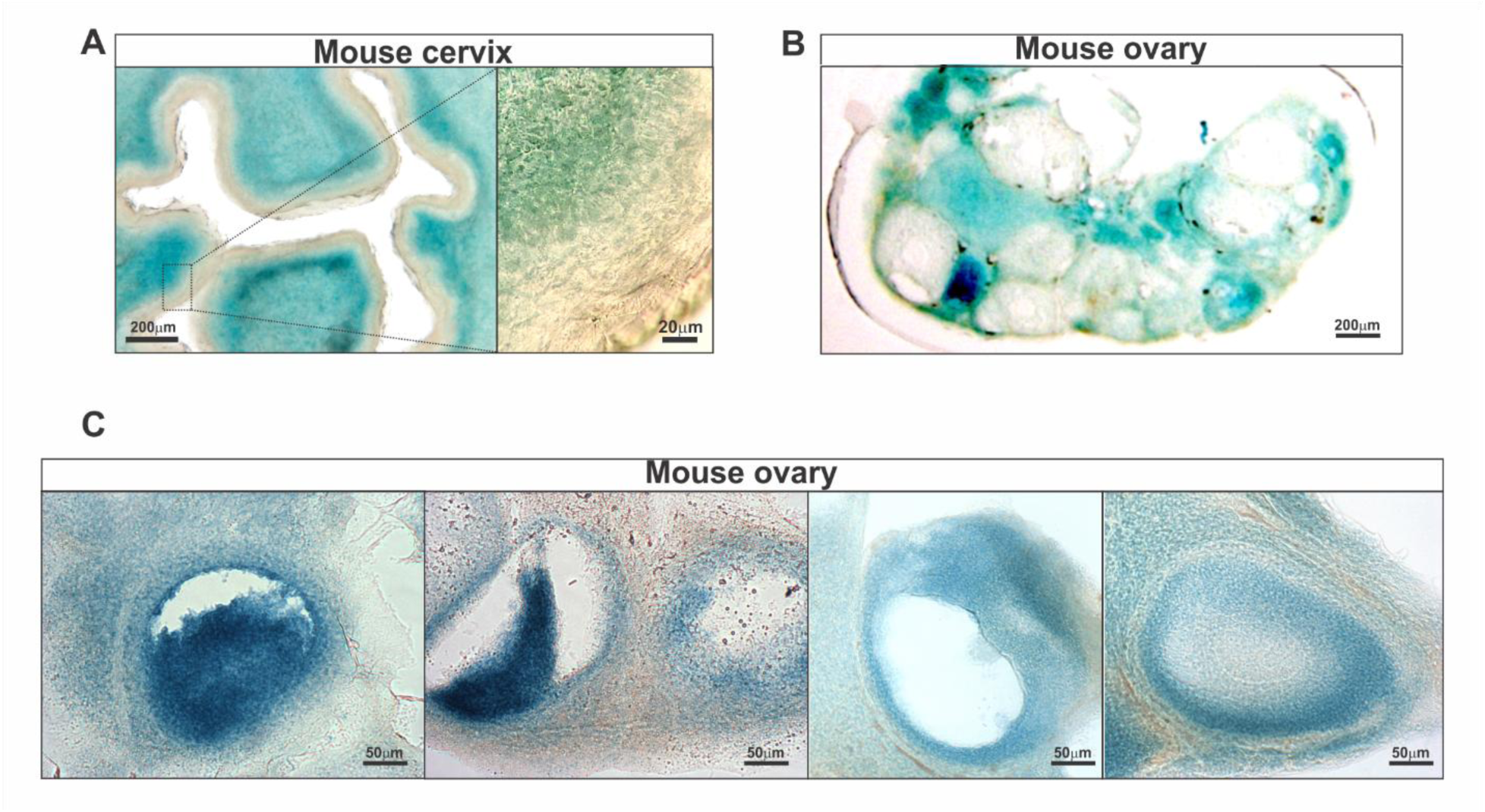
Spag17 expression in mouse cervical and ovarian tissues. Tissues were collected from heterozygous young Spag17/Sox2-Cre female mice (n=3) and stained using a β-Gal Reporter Gene Staining kit. (**A**) Representative images showing *Spag17* expression in the cervix. Note the strong expression in the stromal area. Scale bars, 200 µm and 20 µm. (**B**) Representative image showing *Spag17* expression in the ovary. Scale bar 200 µm. (**C**) Representative images showing expression in granulosa cells, theca cells, and stromal cells. Scale bars 50 µm.

**Figure S2:**
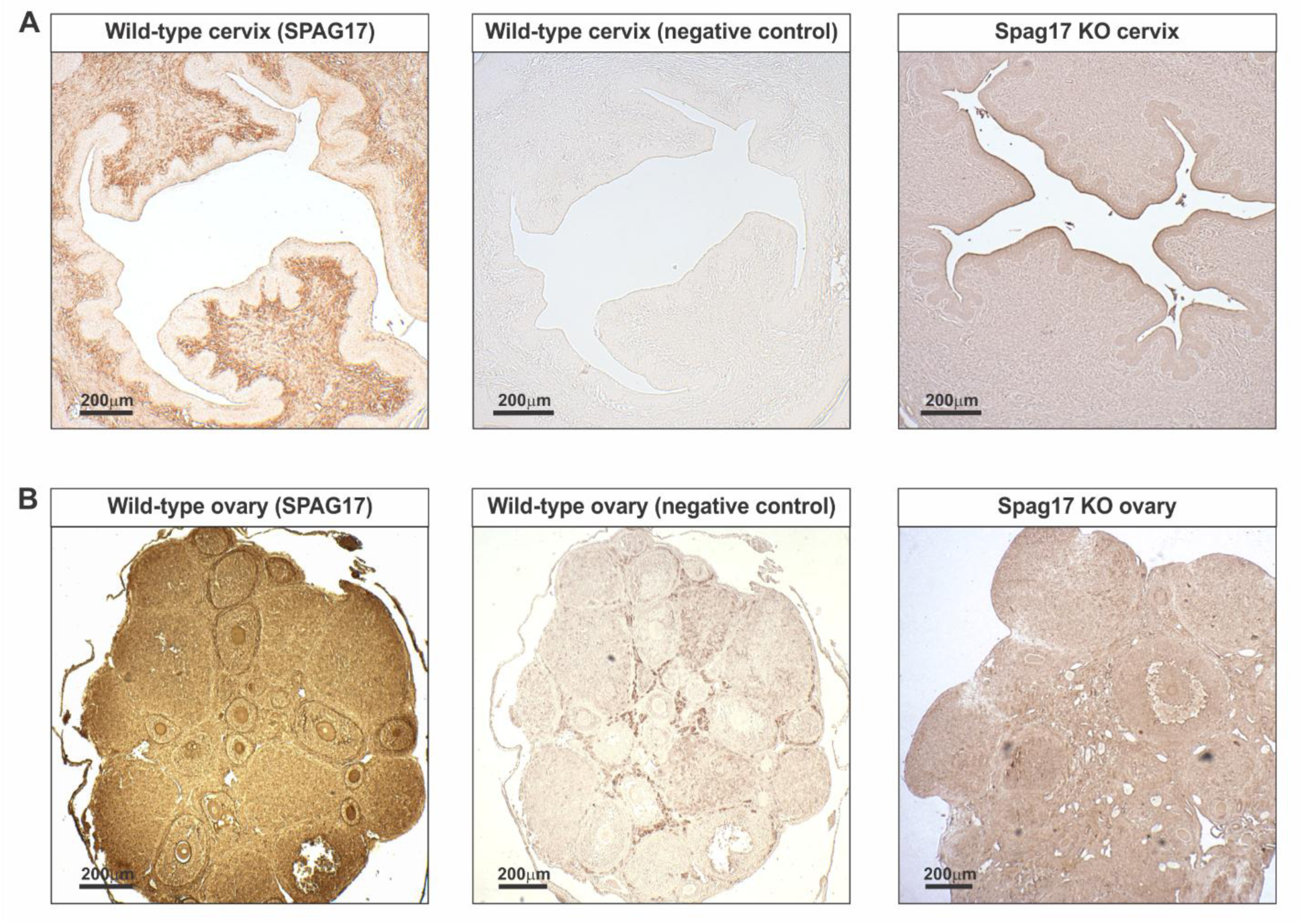
Detection of SPAG17 expression by immunohistochemistry. (**A**) Representative images showing SPAG17 immunolabeling in wild-type cervix, wild-type negative control, and *Spag17* knockout cervix, n=6. (**B**) Representative images showing SPAG17 immunolabeling in wild-type ovary, negative control in wild-type ovary, and *Spag17* knockout ovary, n=6. Scale bars, 200 µm.

**Figure S3:**
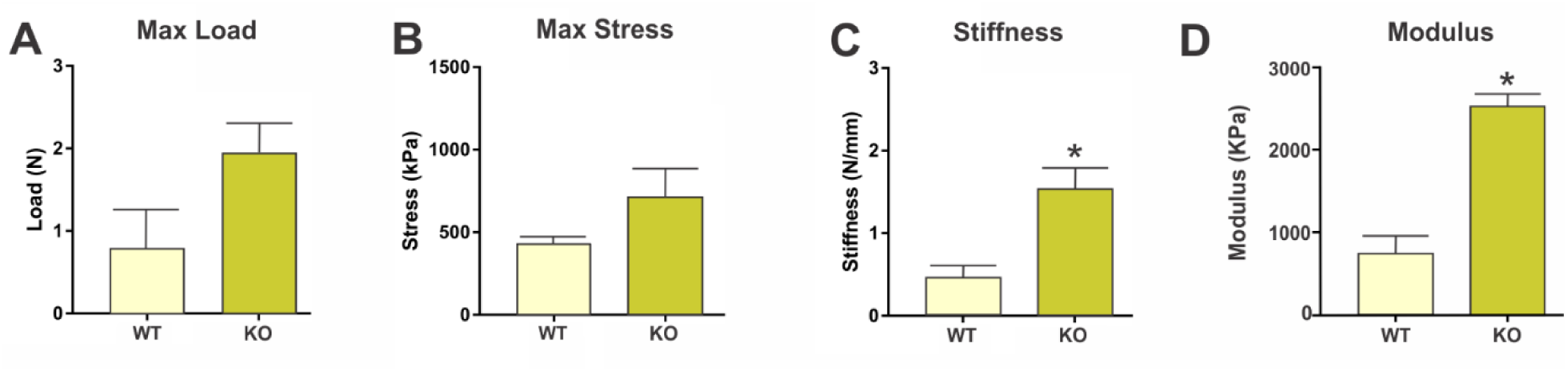
Loss of Spag17 results in increased cervical tissue stiffness. Cervices from two reproductive age females from wild-type (WT) and *Spag17* knockout (KO) mouse line were collected and analyzed using an Instron 5848 testing system. Biomechanical properties were measured in cervices including (**A**) Max load; (**B**) Max stress; (**C**) Stiffness; (**D**) Modulus. Note increased stiffness in the KO tissues. Data are means ± SEM *p<0.05.

**Figure S4:**
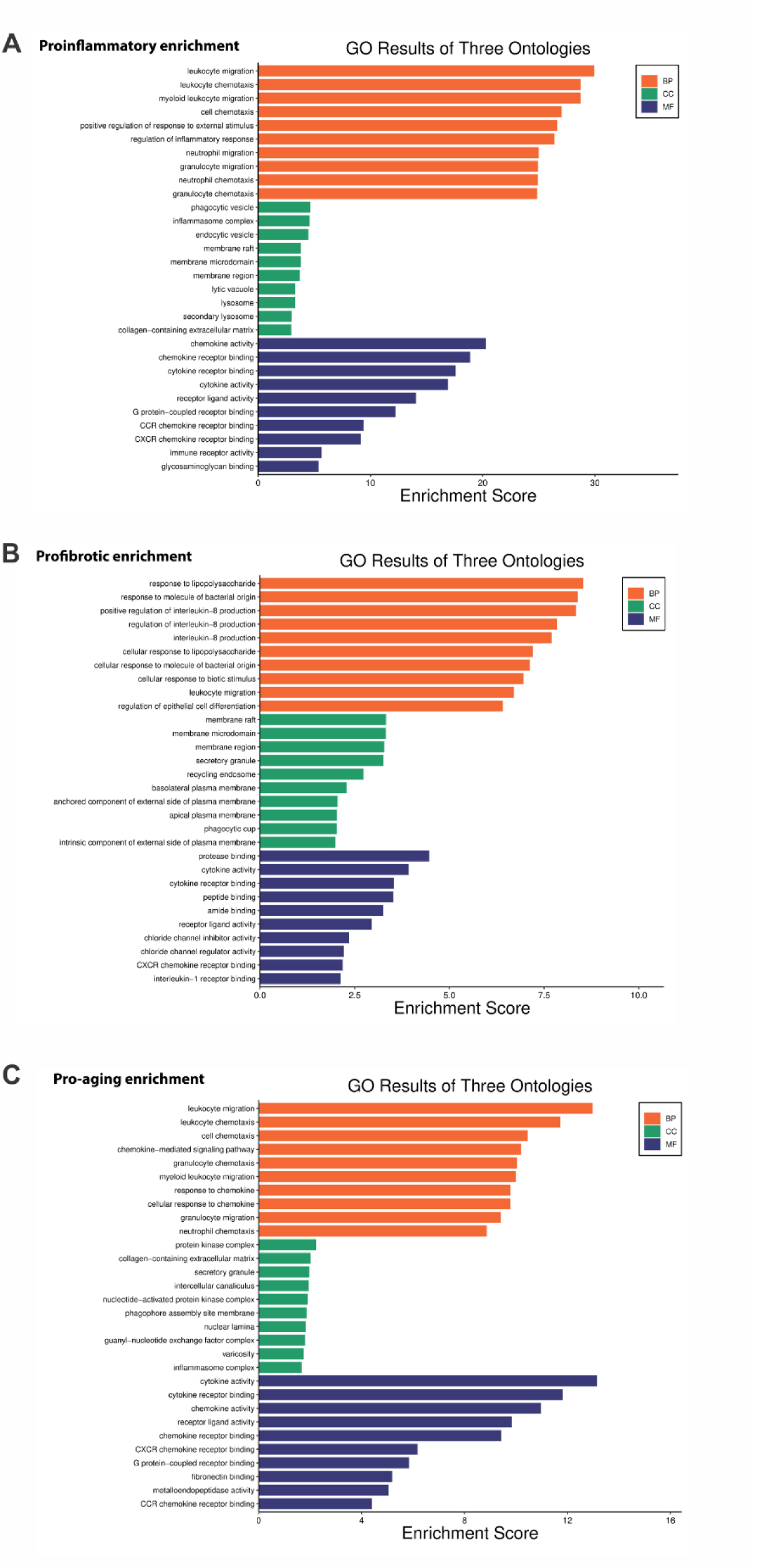
Gene ontologies of biological processes, cellular compartments, and molecular functions in Spag17 knockout vs. wild-type cervices. GO analysis shows increased proinflammatory (**A**), profibrotic (**B**), and aging/senescence (**C**) enrichments in the *Spag17* knockout cervices.

**Figure S5:**
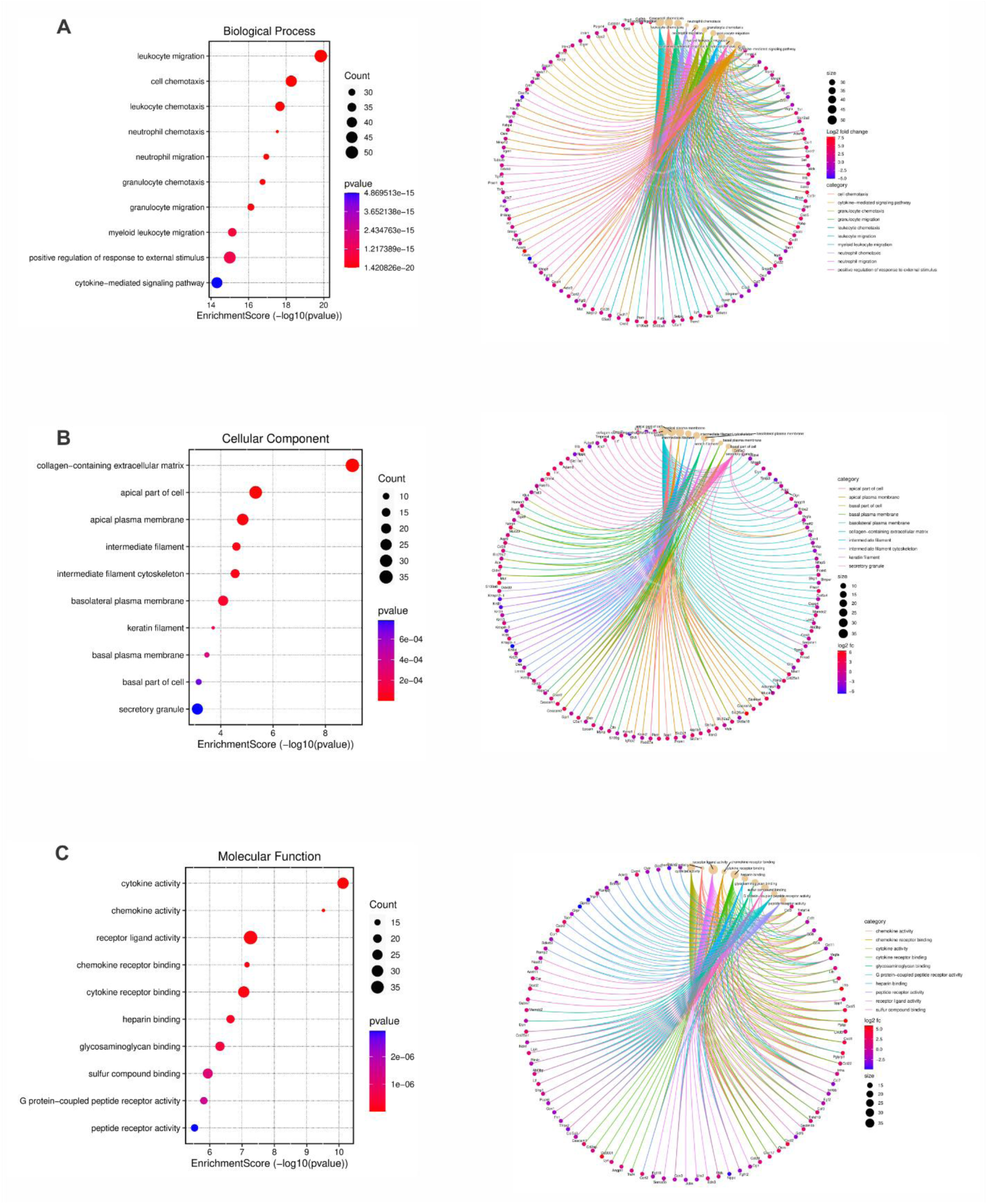
Bubble plot and chord diagrams illustrating RNAseq analysis. Results for enrichment of biological processes (**A**), cellular components (**B**), and molecular functions (**C**) in *Spag17* knockout vs. wild-type mouse cervices.

**Figure S6:**
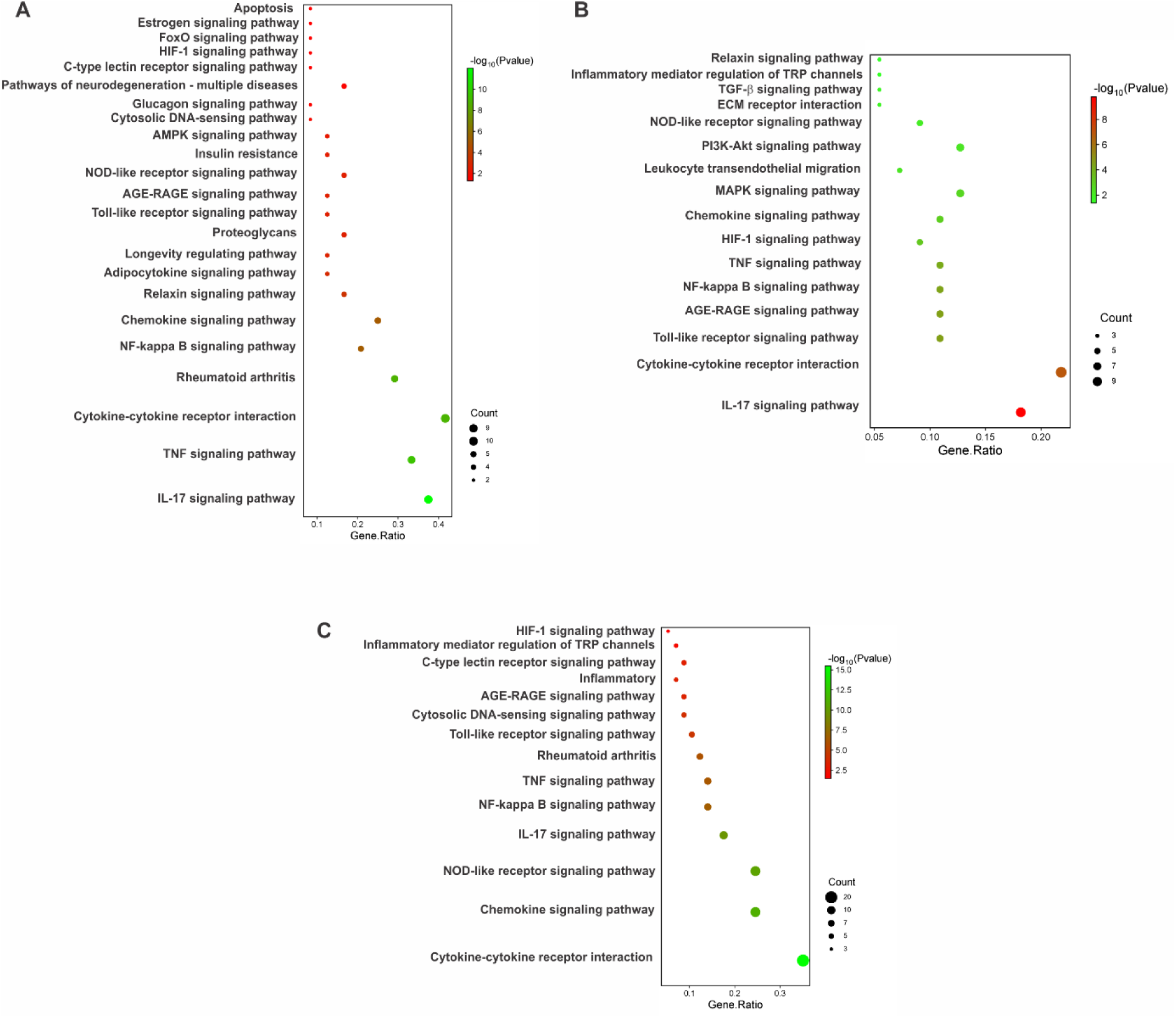
Bubble plots illustrating RNAseq analysis. Results for enrichment of biological processes (**A**), cellular components (**B**), and molecular functions (**C**) in *Spag17* knockout vs. wild-type mouse cervices.

**Figure S7:**
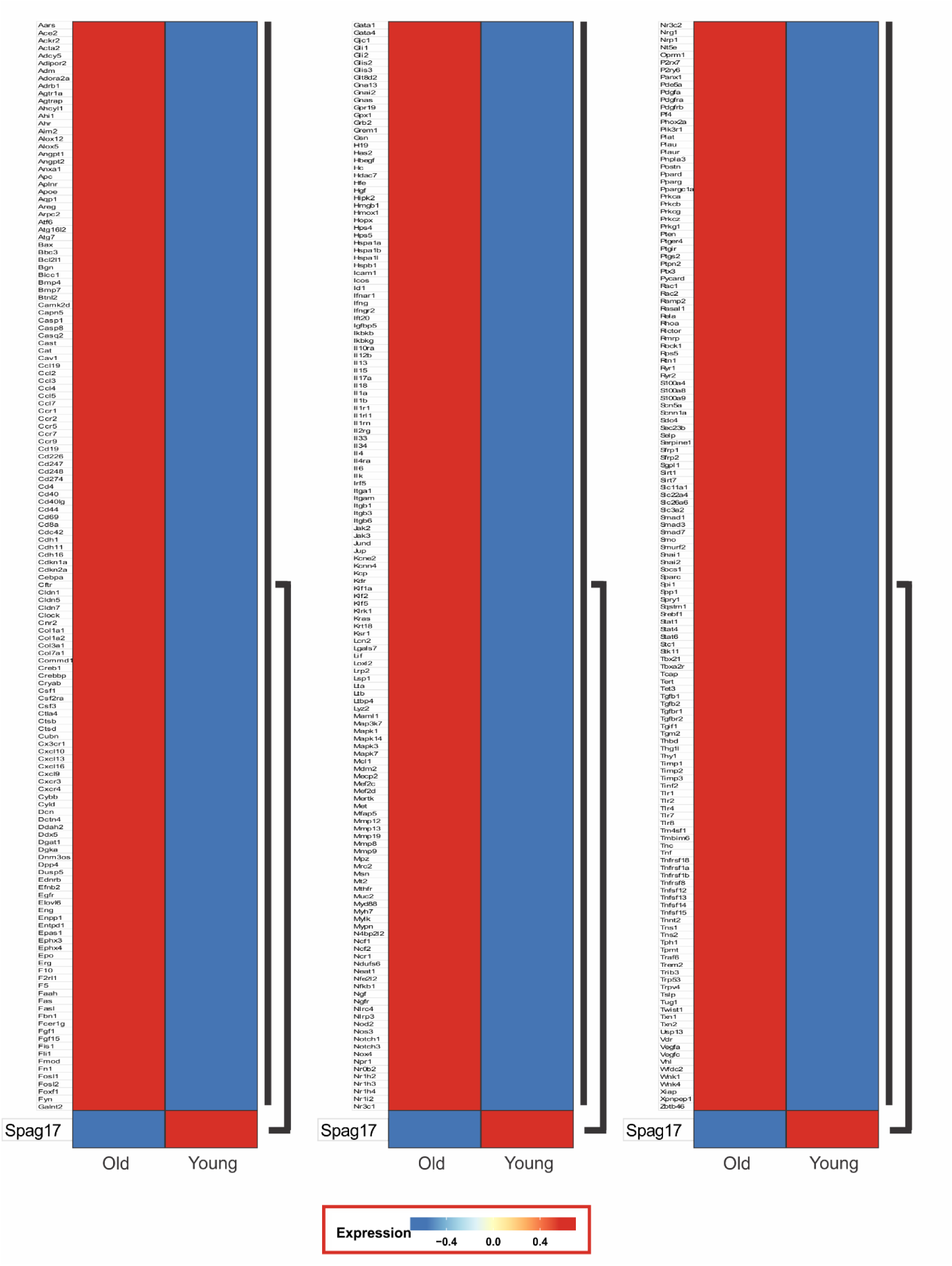
Downregulation of Spag17 in physiological mouse ovarian aging is associated with increased expression of profibrotic genes. scRNAseq data from mouse ovarian samples comparing wild-type young and aged females were obtained from Isola et al., 2024. Note that aged ovaries are characterized by reduced *Spag17* expression and increased expression of fibrosis-associated genes.

**Figure S8:**
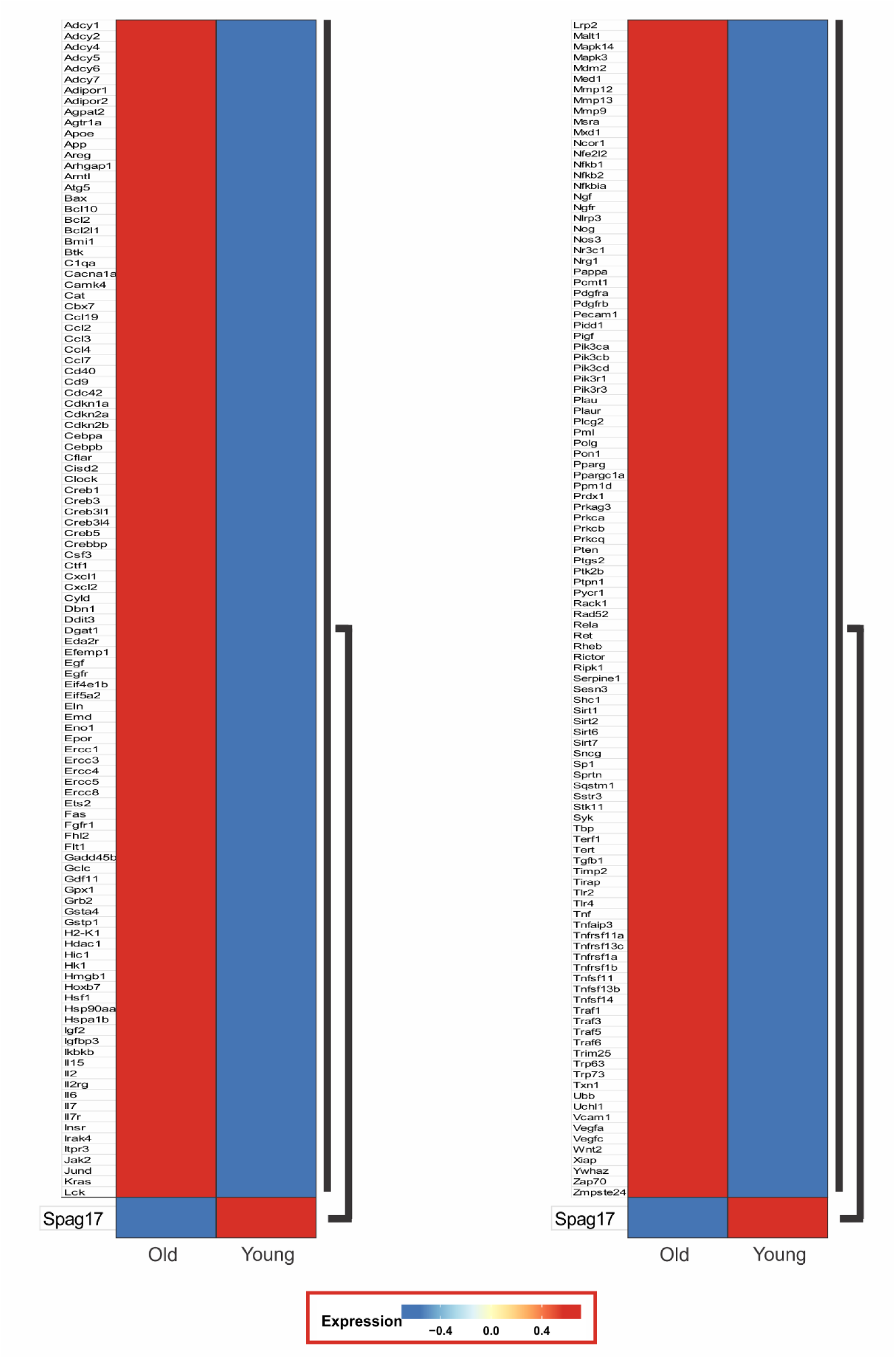
Downregulation of Spag17 in physiological mouse ovarian aging is associated with increased expression of proinflammatory genes. scRNAseq data from mouse ovarian samples comparing wild-type young and aged females were obtained from Isola et al., 2024. Note that aged ovaries are characterized by reduced *Spag17* expression and increased expression of proinflammatory genes.

**Figure S9:**
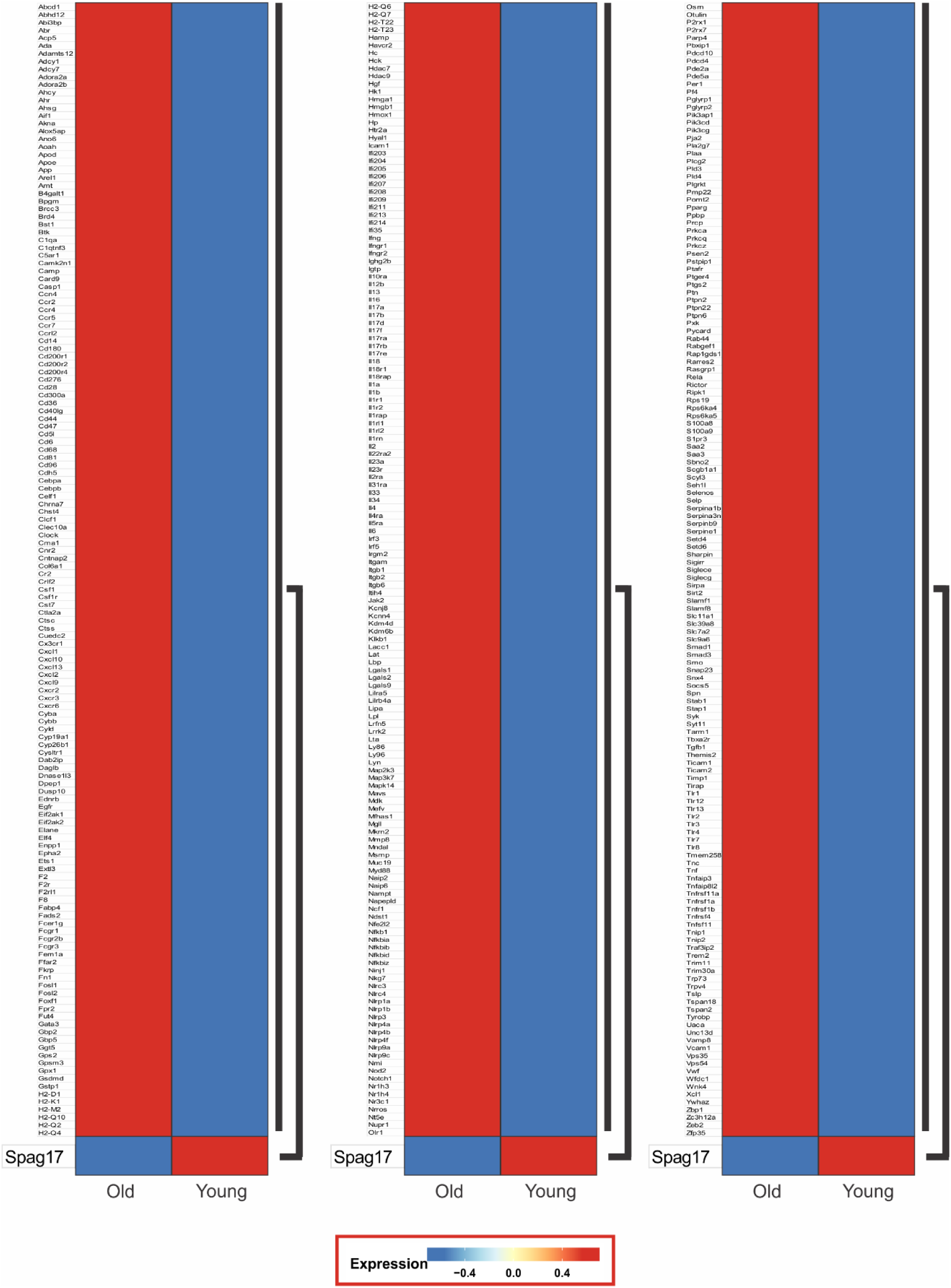
Downregulation of Spag17 in physiological mouse ovarian aging is associated with increased expression of aging/senescence-related genes. scRNAseq data from mouse ovarian samples comparing wild-type young and aged females were obtained from Isola et al., 2024. Note that aged ovaries are characterized by reduced *Spag17* expression and increased expression of aging/senescence-associated genes.

**Figure S10:**
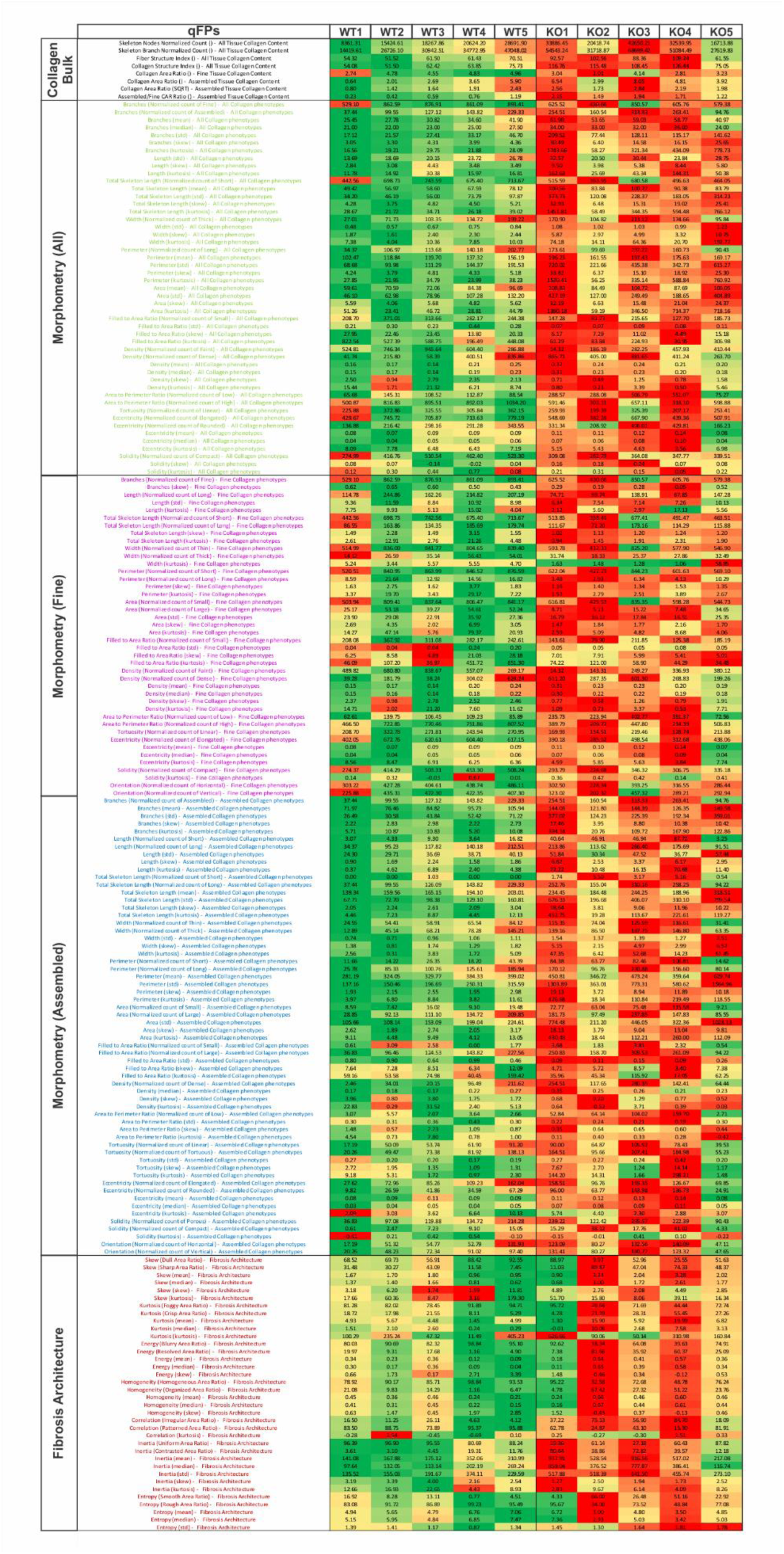
Phenotypic heatmap describing each ovary sample phenotype. The major quantitative fibrosis traits (qFTs) of each ovary sample phenotype are shown as a heatmap (195 in total). Each qFT results from AI-based quantification of PSR collagen histological staining, accounting for the changes and distortions in the fibrosis phenotype for each sample (n=5).

**Figure S11:**
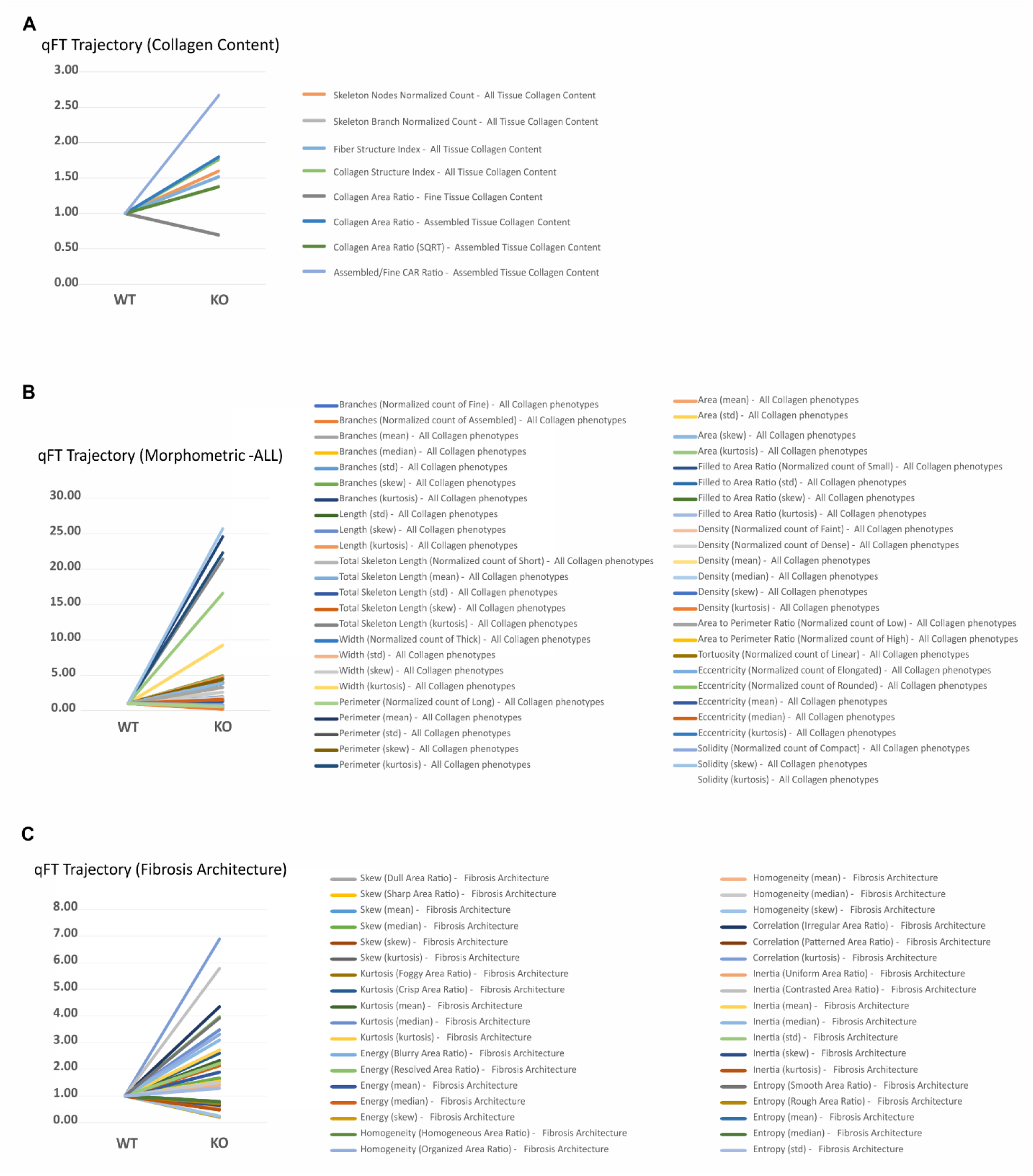
Normalized quantitative fibrosis traits (qFTs) trajectories for each major category. Each qFT from AI-based quantification of PSR collagen histological staining was normalized to the wild-type control, and trajectories were calculated for *Spag17* knockout samples. Graphs represent the trajectories for (**A**) Collagen content, (**B**) Morphometric-All fibers, (**C**) Fibrosis architecture.

**Table S1:** List of profibrotic genes differentially expressed in Spag17 knockout cervices. Gene set enrichment and gene ontology analysis show differences in genes associated with fibrosis.

| Name | log2FoldChange_shrunken | lfcSE_shrunken | p value | padj |
| --- | --- | --- | --- | --- |
| Angpt1 | 0.85943844 | 0.53784574 | 0.000838692 | 0.025900767 |
| Bdkrb1 | -2.934158137 | 0.839806829 | 3.13106E-06 | 0.000443948 |
| Bdkrb2 | -1.855838297 | 0.540562365 | 1.64398E-05 | 0.00149804 |
| Ccl3 | 1.411897189 | 1.198185054 | 0.001225849 | 0.033303663 |
| Ccl4 | 2.165817008 | 1.526502696 | 0.000365565 | 0.014886989 |
| Ccl7 | -2.753211458 | 0.795207866 | 4.36315E-06 | 0.000561286 |
| Ccr1 | 3.338221698 | 1.119042631 | 8.9618E-06 | 0.000977995 |
| Cd14 | 2.952585214 | 1.016860393 | 1.4047E-05 | 0.001364319 |
| Cftr | 1.487401121 | 0.701633349 | 0.000253372 | 0.011690604 |
| Cldn7 | 1.728036936 | 0.523633239 | 2.37304E-05 | 0.001930867 |
| Col25a1 | 2.630192004 | 0.711484747 | 2.68266E-06 | 0.000397792 |
| Cxcl5 | 3.710940613 | 1.663587868 | 3.65228E-05 | 0.002728672 |
| Cxcr4 | 2.663512242 | 0.929504226 | 2.03006E-05 | 0.001705829 |
| Cybb | 1.261246285 | 0.934101248 | 0.001203562 | 0.033021447 |
| Dcn | -0.653389958 | 0.266926439 | 4.23083E-05 | 0.003055165 |
| Fmod | 1.468723422 | 0.96177667 | 0.00071083 | 0.02336357 |
| Fn1 | -1.642568762 | 0.536223281 | 3.99266E-05 | 0.00290477 |
| Gstm6 | -1.741103049 | 0.889718149 | 0.000270253 | 0.012294289 |
| Hmox1 | -1.374151679 | 0.709372091 | 0.000376694 | 0.015184092 |
| Hspa1a | -0.915022873 | 0.431178886 | 0.000195333 | 0.009829897 |
| Il1b | 5.793690138 | 1.247816183 | 7.3921E-08 | 2.6591E-05 |
| Ltb | 1.575204578 | 0.79961739 | 0.000310817 | 0.013357559 |
| Met | 0.937105827 | 0.246784982 | 1.1936E-06 | 0.000214682 |
| Mmp3 | 1.452538201 | 0.828317153 | 0.00049967 | 0.018374674 |
| Mmp9 | 1.512637569 | 1.069017421 | 0.000790545 | 0.024957884 |
| Nlrp12 | 3.035558243 | 1.51085822 | 7.47465E-05 | 0.004823759 |
| Nlrp3 | 1.21930548 | 1.525089645 | 0.002286369 | 0.048568672 |
| S100a8 | 5.629552758 | 1.240528612 | 1.04926E-07 | 3.45456E-05 |
| S100a9 | 5.824525597 | 1.278435009 | 1.02434E-07 | 3.45086E-05 |
| Serpine1 | -1.780600447 | 0.347350056 | 5.67141E-08 | 2.07862E-05 |
| Spp1 | 2.264171165 | 0.926604746 | 6.95856E-05 | 0.004551177 |
| Tnc | -3.463974834 | 0.753519037 | 2.11378E-08 | 9.12448E-06 |
| Tnf | 2.61761883 | 1.288331638 | 9.66799E-05 | 0.005832319 |
| Tnfrsf1b | 1.128720526 | 0.848636285 | 0.001450524 | 0.036928473 |
| Tnfsf14 | 1.61933994 | 1.189941249 | 0.000736327 | 0.02403891 |
| Vegfa | -0.781445914 | 0.352629058 | 0.000115517 | 0.006613766 |

**Table S2:**
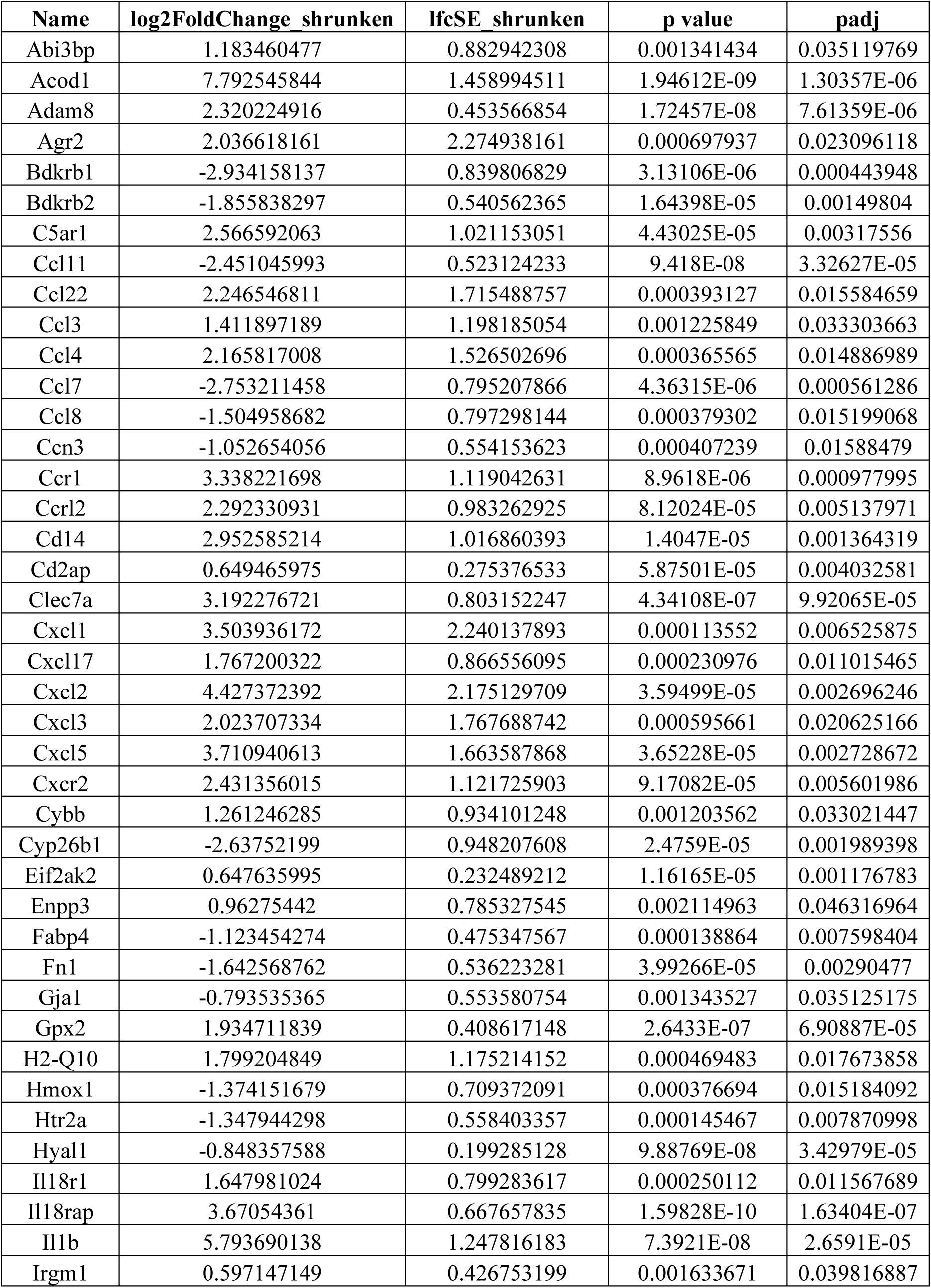
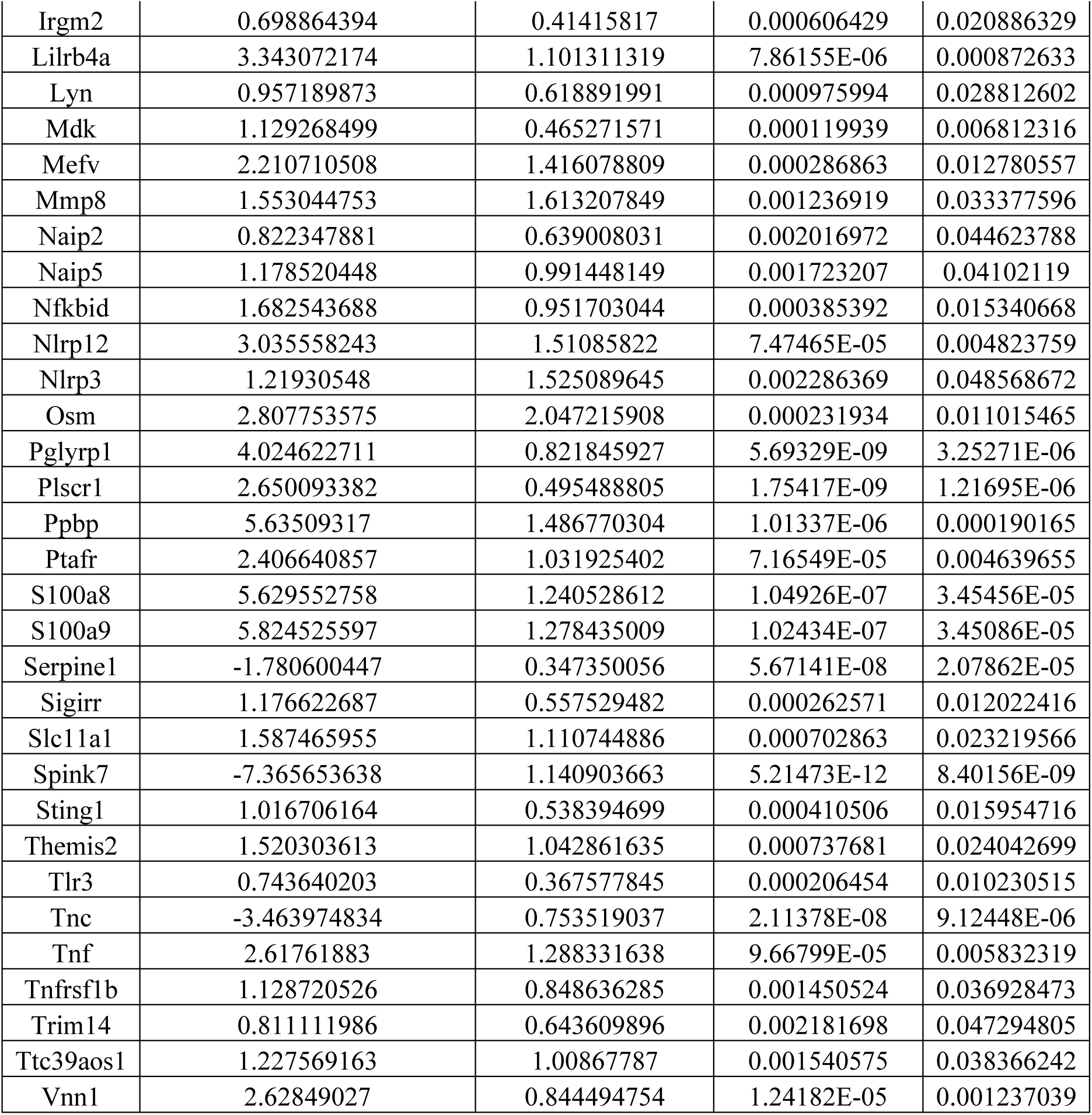
List of proinflammatory genes differentially expressed in Spag17 knockout cervices. Gene set enrichment and gene ontology analysis show differences in genes associated with inflammation.

**Table S3:**
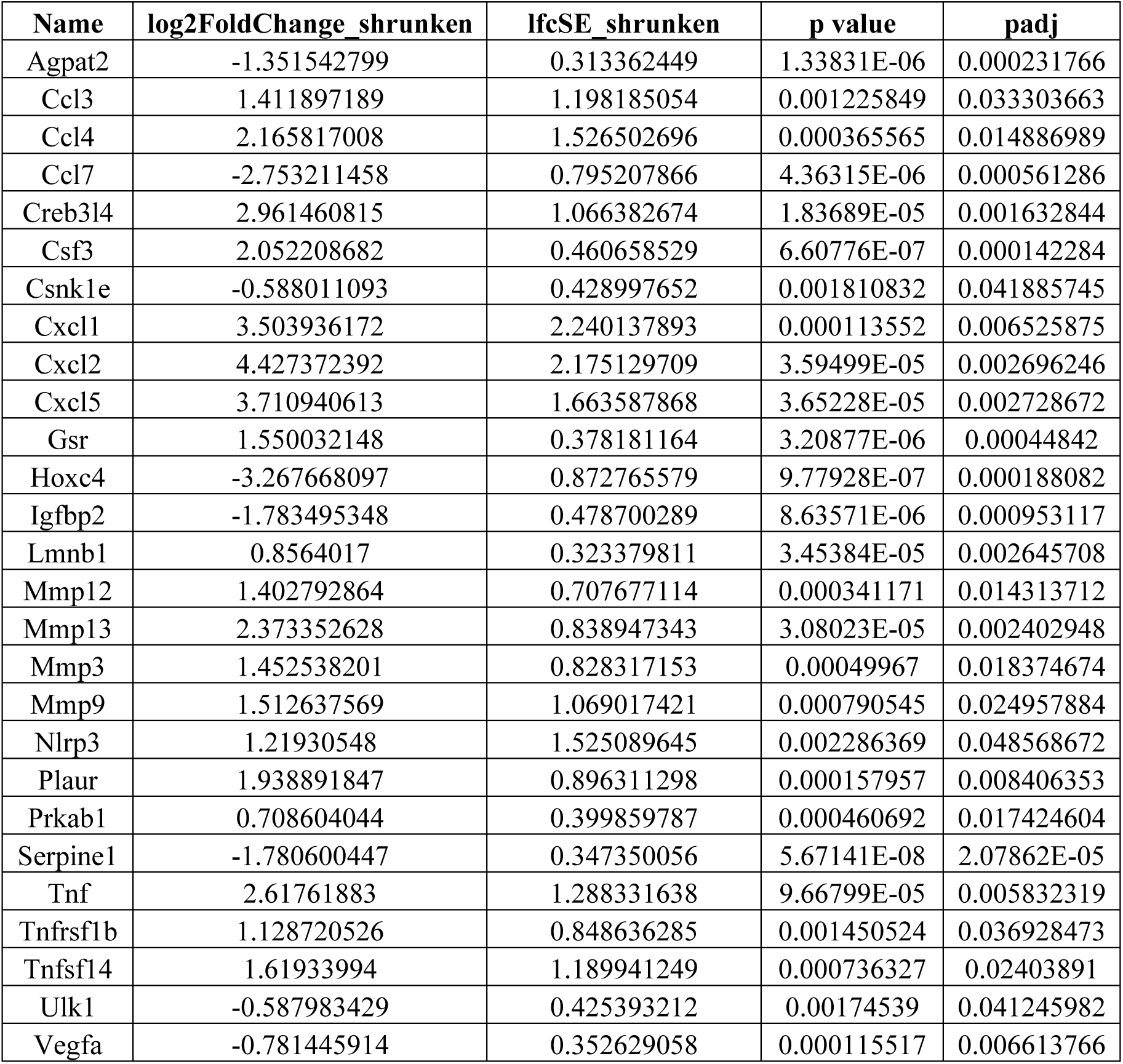
List of senescence-associated genes differentially expressed in Spag17 knockout cervices. Gene set enrichment and gene ontology analysis show differences in genes associated with aging/senescence.

## Notes

### Competing Interest Statement

The authors have declared no competing interest.

## REFERENCES

C. Agudo-Rios, A. Rogers, I. King, V. Bhagat, L.M.T. Nguyen, C. Córdova-Fletes, D. Krapf, J.F. Strauss 3rd, L. Arévalo, G.E. Merges, H. Schorle, E.R.S. Roldan, M.E. Teves. SPAG17 mediates nuclear translocation of protamines during spermiogenesis. Front Cell Dev Biol. (2023).

J. P. O. D. Almeida, T. P. Ribeiro, I. A. D. Medeiros, Aging: molecular pathways and implications on the cardiovascular system. Oxidative Medicine and Cellular Longevity. 2017, 7941563 (2017).

F. Amargant, S. L. Manuel, Q. Tu, W. S. Parkes, F. Rivas, L. T. Zhou, J. E. Rowley, C. E. Villanueva, J. E. Hornick, G. S. Shekhawat, J. J. Wei, M. E. Pavone, A. R. Hall, M. T. Pritchard, F. E. Duncan, Ovarian stiffness increases with age in the mammalian ovary and depends on collagen and hyaluronan matrices. Aging Cell. 19, e13259 (2020).

S. Andrews, FastQC: A Quality Control Tool for High Throughput Sequence Data [Online]. Babraham Bioinformatics (2010).

J.L. Balough, S.S. Dipali, K. Velez, T.R. Kumar, F.E. Duncan, Hallmarks of female reproductive aging in physiologic aging mice. Nat Aging. 12, 1711–1730 (2024).

E. Barnum, J. L. Fey, S.N. Weiss, G. Barila, A.G. Brown, B.K. Connizo, Tensile mechanical properties and dynamic collagen fiber re-alignment of the murine cervix are dramatically altered throughout pregnancy. Journal of Biomechanical Engineering, 139, 061008 (2017).

J.S. Bredfeldt, Y. Liu, M.W. Conklin, P.J. Keely, T.R. Mackie, K.W. Eliceiri, Automated quantification of aligned collagen for human breast carcinoma prognosis. J. Pathol. Inform. 1, 28 (2014).

F. Briand, J. Maupoint, E. Brousseau, N. Breyner, M. Bouchet, C. Costard, T. Leste-Lasserre, M. Petitjean, L. Chen, A. Chabrat, V. Richard, R. Burcelin, C. Dubroca, T. Sulpice, Elafibranor improves diet-induced nonalcoholic steatohepatitis associated with heart failure with preserved ejection fraction in Golden Syrian hamsters. Metabolism, 117, 154707 (2021).

S.M. Briley, S. Jasti, J.M. McCracken, J.E. Hornick, B. Fegley, M.T. Pritchard, F.E. Duncan, Reproductive age-associated fibrosis in the stroma of the mammalian ovary. Reproduction, 152, 245–260 (2016).

A.R. Davalos, JP. Coppe, J. Campisi, PY. Desprez, Senescent cells as a source of inflammatory factors for tumor progression. Cancer Metastasis Rev, 29, 273–283 (2010).

S.S. Dipali, C.D. King, J.P. Rose, J.E. Burdette, J. Campisi, B. Schilling, F.E. Duncan, Proteomic quantification of native and ECM-enriched mouse ovaries reveals an age-dependent fibro-inflammatory signature. Aging, 15, 10821–10855 (2023).

A. Dobin, C.A. Davis, F. Schlesinger, J. Drenkow, C. Zaleski, S. Jha, P. Batut, M. Chaisson, T.R. Gingeras, STAR: ultrafast universal RNA-seq aligner. Bioinformatics, 29, 15–21 (2013).

F.E. Duncan, R. Confino, M.E. Pavone, Chapter 9 - Female reproductive aging: From consequences to mechanisms, markers, and treatments. Conn’s Handbook of Models for Human Aging (Second Edition), 109–130 (2018).

P. Ewels, M. Magnusson, S. Lundin, M. Käller, MultiQC: summarize analysis results for multiple tools and samples in a single report. Bioinformatics, 32, 3047–3048 (2016).

K.G. Foley, M.T. Pritchard, F.E. Duncan, Macrophage-derived multinucleated giant cells: hallmarks of the aging ovary. Reproduction, 161 V5–V9 (2021).

J. Guo, X. Huang, L. Duo, M. Yan, T. Shen, W. Tang, J. Li, Aging and aging-related diseases: from molecular mechanisms to interventions and treatments. Signal Transduction and Target Therapy, 7, 391 (2022).

J. Harrow, A. Frankish, J.M. Gonzalez, E. Tapanari, M. Diekhans, F. Kokocinski, B.L. Aken, D. Barrell, A. Zadissa, S. Searle, I. Barnes, A. Bignell, V. Boychenko, T. Hunt, M. Kay, G. Mukherjee, J. Rajan, G. Descaio-Reyes, G. Saunders, C. Saunders, C. Steward, R. Harte, M. Lin, C. Howald, A. Tanzer, T. Derrien, J. CHrast, N. Walters, S. Balasubramanian, B. Pei, M. Tress, J.M. Rodriguez, I. Ezkurdia, J. van Baren, M. Brent, D. Haussler, M. Kellis, A. Valencia, A. Reymond, M. Gerstein, R. Guidό, T.J. Hubbard, GENCODE: the reference human genome annotation for The ENCODE Project. Genome Research, 22, 1760–1774 (2012).

S. Hartley, J.C. Mullikin, QoRTs: a comprehensive toolset for quality control and data processing of RNA-Seq experiments. BMC Bioinformatics, 16, 224 (2015).

K. Inada, S. Hayashi, T. Iguchi, T. Sato. Establishment of a primary culture model of mouse uterine and vaginal stroma for studying in vitro estrogen effects. Exp Biol Med, 231,303–10 (2006).

J.A. Inia, G. Stokman, M.C. Morrison, N. Worms, L. Verschuren, M.P.M. Caspers, A.L. Menke. L. Petitjean, L. Chen, M. Petitjean, J.W. Jukema, H.M.G. Princen, A.M. van den Hoek, Semaglutide Has Beneficial Effects on Non-Alcoholic Steatohepatitis in Ldlr-/-.Leiden Mice. International Journal of Molecular Sciences, 24, 8494 (2023).

J.V.V. Isola, S.R. Ocañas, C.R. Hubbart, S. Ko, S.A. Mondal, J.D. Hense, H.N.C. Carter, A. Schneider, S. Kovats, J. Alberola-Ila, W.M. Freeman, M.B. Stout. A single-cell atlas of the aging mouse ovary. Nat Aging, 1,145–162 (2024).

A.S.K. Jones, D. F. Hannum, J.H. Machlin, A. Tan, Q. Ma, N.D. Ulrich, YC. Shen, M. Ciarelly, V. Padmanabhan, E.E. Marsh, S. Hammoud, J.Z. Li, A. Shikanov, Cellular atlas of the human ovary using morphologically guided spatial transcriptomics and single-cell sequencing. Science Advances, 10, eadm7506 (2024).

E. Kazarian, H. Y. Son, P. Sapao, W. Li, Z. Zhang, J.F. Strauss, M.E. Teves, SPAG17 is required for male germ cell differentiation and fertility. International Journal of Molecular Sciences, 19, 1252 (2018).

R. Kolde, Pheatmap: pretty heatmaps [Online]. R Package Version 1.0.10. (2012).

R. Kostadinova, S. Strӧbel, L. Chen, K. Fiaschetti-Egli, J. Gadient, A. Pawlowska, L. Petitjean, M. Bieri, E. Thoma, M. Petitjean, Digital pathology with artificial intelligence analysis provides insight to the efficacy of anti-fibrotic compounds in human 3D MASH model. Science Reports, 14, 5885 (2024).

M. Laopaiboon, P. Lumbiganon, N. Intarut, R. Mori, T. Ganchimeg, J.P. Vogel, J.P. Souza, A.M. Gülmezoglu, WHO Multicountry Survey on Maternal Newborn Health Research Network, Advanced maternal age and pregnancy outcomes: a multicountry assessment. BJOG, 121, 49–56 (2014).

R. Li, K. Hu, H. Liu, M.R. Green, L.J. Zhu, OneStopRNAseq: a web application for comprehensive and efficient analyses of RNA-Seq data. Genes, 11, 1165 (2020).

Liao Y, Smyth GK, Shi W. featureCounts: an efficient general purpose program for assigning sequence reads to genomic features. Bioinformatics, 30, 923–30 (2014).

Y. Liu, A. Keikhosravi, G.S. Mehta, C.R. Drifka, K.W. Eliceiri, Methods for quantifying fibriliar collagen alignment. Methods Mol Biol, 1627, 429–451 (2017).

M.I. Love, W. Huber, S. Anders, Moderated estimation of fold change and dispersion for RNA-seq data with DESeq2. Genome Biology, 15, 550 (2014).

Luo W, Brouwer C. Pathview: an R/Bioconductor package for pathway-based data integration and visualization. Bioinformatics, 29, 1830–1 (2013).

D. Muñoz-Espín, M. Serrano. Cellular senescence: from physiology to pathology. Nat Rev Mol Cell Biol, 7, 482–96 (2014).

Y. Nakamura, H. Miyaaki, S. Miuma, Y. Akazawa, M. Fukusima, R. Sasaki, M. Haraguchi, A. Soyama, M. Hidaka, S. Eguchi, K. Makao, Automated fibrosis phenotyping of liver tissue from non-tumor lesions of patients with and without hepatocellular carcinoma after liver transplantation for non-alcoholic fatty liver disease. Hepatology International, 16, 555–561 (2022).

P. Sapao, B. Shi, E.D.O. Roberson, J. Atkinson, J. Strauss, M. Teves, J. Varga, Reduced SPAG17 expression, links dysfunctional cilia, morphogen signaling activation and multiple organ fibrosis: novel target for systemic sclerosis [abstract]. American College of Rheumatology, 70 (2018)

P. Sapao, E.D.O. Roberson, B. Shi, S. Assassi, B. Skaug, F. Lee, A. Naba, B.E. Perez White, C. Cordova-Fletes, PS. Tsou, A.H. Sawalha, J.E. Gudjonsson, F. Ma, P. Verma, D. Bhattacharyya, M. Carms, J.F. Strauss III, D. Sicard, D.J. Tschumperlin, M.I. Champer, P.J. Campagnola, M.E. Teves, J. Varga, Reduced SPAG17 expression in systemic sclerosis triggers myofibroblast transition and drives fibrosis. The Journal of Investigative Dermatology, 143, 284–293 (2023).

M. Selman, A. Pardo, Fibroageing: An ageing pathological feature driven by dysregulated extracellular matrix-cell mechanobiology. Ageing Research Reviews, 70, 101393 (2021).

S. Shen, J.W. Park, Z. Lu, L. Lin, M.D. Henry, Y.N. Wu, Q. Zhou, Y. Xing, rMATS: robust and flexible detection of differential alternative splicing from replicate RNA-Seq data. Proceedings of the National Academy of Sciences of the United States of America, 111, E5593–E5601 (2014).

G. Soon, A. Wee, Updates in the quantitative assessment of liver fibrosis for nonalcoholic fatty liver disease: Histological perspective. Clinical and Molecular Hepatology, 27, 44–57 (2021).

A. Subramanian, P. Tamayo, V.K. Mootha, S. Mukherjee, B.L. Ebert, M.A. Gillette, A. Paulovich, S.L. Pomeroy, T.R. Golub, E.S. Lander, J.P. Mesirov, Gene set enrichment analysis: a knowledge-based approach for interpreting genome-wide expression profiles. Proceedings of the National Academy of Sciences of the United States of America, 102, 15545–15550 (2005).

V. Suryadevara, A.D. Hudgins, A. Rajesh, A. Pappalardo, A. Karpova, A.K. Dey, A. Hertzel, A. Agudelo, A. Rocha, B. Soygur, B. Schilling, C.M. Carver, C. Aguayo-Mazzucato, D.J. Baker, D.A. Bernlohr, D. Jurk, D.B. Mangarova, E.M. Quardokus, E.A.L. Enninga, E.L. Schmidt, F. Chen, F.E. Duncan, F. Cambuli, G. Kaur, G.A. Kuchel, G. Lee, H.E. Daldrup-Link, H. Martini, H. Phatnani, I.M. Al-Naggar, I. Rahman, J. Nie, J.F. Passos, J.C. Silverstein, J. Campisi, J. Wang, K. Iwasaki, K. Barbosa, K. Metis, K. Nernekli, L.J. Niedernhofer, L. Ding, L. Wang, L.C. Adams, L. Ruiyang, M.L. Doolittle, M.G. Teneche, M.J. Schafer, M. Xu, M. Hajipour, M. Boroumand, N. Basisty, N. Sloan, N. Slavov, O. Kuksenko, P. Robson, P.T. Gomez, P. Vasilikos, P.D. Adams, P. Carapeto, Q. Zhu, R. Ramasamy, R. Perez-Lorenzo, R. Fan, R. Dong, R.R. Montgomery, S. Shaikh, S. Vickoviic, S. Yin, S. Kang, S. Suvakov, S. Khosla, V.D. Garovic, V. Menon, Y. Xu, Y. Song, Y. Suh, Z. Dou, N. Neretti, SenNet recommendations for detecting senescent cells in different tissues. Nature Reviews Molecular Cell Biology, 12,1001–1023 (2024).

T. Tchkonia, Y. Zhu, J. van Deursen, J. Campisi, J.L. Kirkland. Cellular senescence and the senescent secretory phenotype: therapeutic opportunities. J Clin Invest, 3, 966–72 (2013).

M.E. Teves, G. Sundaresan, D.J. Cohen, S.L. Hyzy, I. Kajan, M. Maczis, Z. Zhang, R.M. Costanzo, J. Zweit, Z. Schwartz, B.D. Boyan, J.F. Strauss III, Spag17 deficiency results in skeletal malformations and bone abnormalities, PLoS One, 10, e0125936 (2015)

M.E. Teves, Z, Zhang, R.M. Costanzo, S.C. Henderson, F.D. Corwin, J. Zweit, G. Sundaresan, M. Subler, F.N. Salloum, B.K. Rubin, J.F. Strauss III, Sperm-associated antigen-17 gene is essential for motile cilia function and neonatal survival. American Journal of Respiratory Cell and Molecular Biology, 48, 765–772 (2013)

T. Umehara, J.S. Richards, M. Shimada, The stromal fibrosis in aging ovary. Aging, 10, 9–10 (2018).

S. Wang, K. Li, E. Pickholz, R. Dobie, K.P. Matchett, N.C. Henderson, C. Carrico, I. Driver, M.B. Jensen, L. Chen, M. Petitjean, D. Bhattacharya, M.I. Fiel, X. Liu, T. Kisseleva, U. Alon, M. Adler, R. Medzhitov, S.L. Friedman, An autocrine signaling circuit in hepatic stellate cells underlies advanced fibrosis in nonalcoholic steatohepatitis. Science Translational Medicine, 15, eadd3949 (2023).

I. Winkler, A. Tolkachov, F. Lammers, P. Lacour, K. Daugelaite, N. Schneider, ML. Koch, J. Panten, F. Grünschläger, T. Poth, B. Machado de Ávila, A. Schneider, S. Haas, D.T. Odom, A. Gonçalves, The cycling and aging mouse female reproductive tract at single-cell resolution. Cell, 187, 981–998 (2024).

Y. Wu, M. Li, J. Zhang, S. Wang, Unveiling uterine aging: Much more to learn. Ageing Research Reviews, 86, 101879 (2023).

M. Wu, W. Tang, Y. Chen, L. Xue, J. Dai, Y. Li, X. Zhu, C. Wu, J. Xiong, J. Zhang, T. Wu, S. Zhou, D. Chen, C. Sun, J. Yu, H. Li, Y. Guo, Y. Huang, Q. Zhu, S. Wei, Z. Zhou, M. Wu, Y. Li, T. Xiang, H. Qiao, S. Wang. Spatiotemporal transcriptomic changes of human ovarian aging and the regulatory role of FOXP1. Nat Aging, 4, 527–545 (2024).

G. Yu, L.G. Wang, Y. Han, Q.Y. He, clusterProfiler: an R package for comparing biological themes among gene clusters. OMICS, 16, 284–287 (2012).

Z. Zhang, F. Schlamp, L. Huang, H. Clark, L. Brayboy, Inflammaging is associated with shifted macrophage ontogeny and polarization in the aging mouse ovary, Reproduction, 159, 325–337 (2020).

C. Zhou, Q. Guo, J. Lin, M. Wang, Z. Zeng, Y. Li, X. Li, Y. Xiang, Q. Liang, J. Liu, T. Wu, Y. Zeng, S. He, S. Wang, H. Zeng, X. Liang. Single-Cell Atlas of Human Ovaries Reveals The Role Of The Pyroptotic Macrophage in Ovarian Aging. Adv Sci (Weinh), (2024).

